# Extensive alternative splicing transitions during postnatal skeletal muscle development are required for calcium handling functions

**DOI:** 10.1101/124230

**Authors:** Amy E. Brinegar, Zheng Xia, James A. Loehr, Wei Li, George G. Rodney, Thomas A. Cooper

**Affiliations:** Department of Molecular & Cellular Biology, Baylor College of Medicine, Houston, TX 77030; Division of Biostatistics, Dan L Duncan Cancer Center, Baylor College of Medicine, Houston, TX 77030; Department of Molecular Physiology & Biophysics, Baylor College of Medicine, Houston, TX 77030; Department of Pathology & Immunology, Baylor College of Medicine, Houston, TX 77030

## Abstract

Postnatal development of skeletal muscle is a highly dynamic period of tissue remodeling. Here we used RNA-seq to identify transcriptome changes from late embryonic to adult mouse muscle and demonstrate that alternative splicing developmental transitions impact muscle physiology. The first two weeks after birth are particularly dynamic for differential gene expression and AS transitions, and calcium-handling functions are significantly enriched among genes that undergo alternative splicing. We focused on the postnatal splicing transitions of three calcineurin A genes, calcium-dependent phosphatases that regulate multiple aspects of muscle biology. Redirected splicing of calcineurin A to the fetal isoforms in adult muscle and in differentiated C2C12 slows the timing of muscle relaxation, promotes nuclear localization of calcineurin targets Nfatc3 and Nfatc2, and affects expression of Nfatc transcription targets. The results demonstrate a previously unknown specificity of calcineurin isoforms as well as the broader impact of AS during muscle postnatal development.

## Introduction

There is a 50-fold increase in body weight during murine postnatal development, 50% of which is contributed by skeletal muscle (1). Skeletal muscle tissue undergoes dynamic remodeling after birth to transition to the functional requirements of adult tissue. While embryonic development of skeletal muscle and regeneration in adult skeletal muscle has been extensively studied, the physiological transitions of postnatal muscle are poorly understood (2-5). In rodents, as in humans, skeletal muscle at birth is immature with low functionality as illustrated by poor mobility of newborns. After a period of active proliferation of myogenic progenitor satellite cells and fusion to form myofibers during late embryonic development, myofiber number remains constant after birth. Postnatal skeletal muscle growth is primarily by myofiber hypertrophy and fusion of proliferating satellite cells that is limited to within the first few weeks after birth (6,7). Satellite cells make up approximately 11% of muscle nuclei at postnatal day 14, but by week 17, the fraction of satellite cell nuclei drops to 3% (6) reflecting their transition from contributing to myofiber growth to quiescent adult muscle stem cells (8). The tibialis anterior muscle cross sectional area increases seven-fold from postnatal day 1 (PN1) to PN28 in mice with a concomitant five-fold increase in maximal isotonic force (9). Importantly, while the increased isotonic force is due primarily to increased muscle size, there is a six-fold increase in intrinsic mechanical function from PN1 to PN28 that is size-independent. The basis for the increased intrinsic function is not completely understood but correlates best with increased myofibril size, a transition of myosin heavy chain isoforms, and changes in metabolism and calcium handling (9). T-tubules and sarcoplasmic reticulum, the cellular structures required for excitation contraction coupling, rapidly mature within the first three weeks after birth (10). The majority of skeletal muscles in mice switch from slow, type I fibers to fast, type II fibers (11). In addition to the differences in metabolism, fast and slow fibers have different cytosolic calcium concentrations of 30 nM and 50-60 nM, respectively (12).

While changes in gene expression are well established mediators of skeletal muscle hypertrophy and atrophy the role of different protein isoforms generated by alternative splicing in muscle physiology has not been extensively explored. We hypothesize that there is a substantial role for protein isoform transitions produced by coordinated alternative splicing in the transition to adult skeletal muscle physiology. Alternative splicing generates proteome diversity including isoforms with tissue specific or developmental stage-specific functions (13-15). Ninety-five percent of human intron-containing genes are alternatively spliced; however, opinions differ with regard to the extent to which this huge diversity is regulated to provide a functional outcome (16-18). Several studies have shown that the majority of alternative splicing is not conserved, however alternative exons that are tissue-specific or regulated during periods of physiological change show high levels of conservation of the variable protein segment, suggesting functional importance; functionality is further supported when the timing of the splicing transition is also conserved (14,19,20). For example, alternative splicing transitions during postnatal development of both brain and heart have been associated with functional consequences (13,21).

While a number of alternative splicing transitions during skeletal muscle postnatal development have been identified, little is known regarding the functional consequences (22-26). Recent reports demonstrate functional consequences using knock out of alternative exons to force expression of inappropriate isoforms of Ca_V_1.1 or titin resulting in reduced contractile force and substantial histopathology, respectively (27,28). Simultaneous reversion to fetal splicing patterns of four vesicular trafficking genes in adult mouse skeletal muscle demonstrated the requirement for the adult isoforms in myofiber structure and physiology (29). Reversion to fetal splicing patterns in skeletal muscle is a hallmark of myotonic dystrophy and results in altered function including myotonia due to failure to express the adult isoform of the CLCN1 mRNA (30,31). The full extent to which alternative splicing contributes to normal postnatal muscle development remains unknown since its role has not been systematically examined on a genome-wide scale.

Among a large number of RNA binding proteins that regulate alternative splicing, the Muscleblind-like (MBNL) and CUG-BP Elav-like family (CELF) families are the best characterized for regulating splicing transitions during postnatal development in heart and skeletal muscle (13,14,32). Disruption of MBNL and CELF RNA processing activities by the repeat containing RNAs expressed from microsatellite expansions cause the pathogenic effects in myotonic dystrophy (33). MBNL and CELF families regulate separate as well as overlapping subsets of alterative splicing events and most often show antagonistic regulation of the shared splicing events (34,35).

We performed a systematic analysis of genome-wide gene expression and alternative splicing transitions in mouse gastrocnemius muscle by RNA-seq of five time points between embryonic day 18.5 (E18.5) and adult. The results show extensive regulation of both gene expression and alternative splicing that is particularly active within the first two weeks after birth. Of the transitions that occur between E18.5 and adult, 55% and 56% of genes that undergo alternative splicing or differential expression, respectively occur between postnatal day 2 (PN2) and PN14. Interestingly, 58% of the splicing transitions that occur between PN2 and PN14 show little change before and after these time points identifying a subset of splicing transitions that are not contiguous with ongoing fetal transitions, but rather are limited to the first two weeks after birth. The genes that undergo differential gene expression and alternative splicing show minimal overlap suggesting independent mechanisms of transcriptional and post-transcriptional regulation. Differentially expressed genes were enriched for mitochondrial functions while genes that undergo alternative splicing transitions were enriched for calcium handling, cell-cell junction, and endocytosis. We show that more than 50% of the splicing transitions tested were conserved between mouse and human with regard to the direction and timing of the transition strongly suggesting functional significance. We used morpholino oligonucleotides to re-direct splicing of all three calcineurin A genes (*Ppp3ca, Ppp3cb*, and/or *Ppp3cc*) to the fetal isoforms in adult mouse flexor digitorum brevis muscle and differentiated C2C12 myotubes. The results demonstrate that the fetal isoforms of calcineurin A specifically cause nuclear localization of Nfatc2 and Nfatc3, but not Nfatc1. These results identify protein isoform transitions that occur during postnatal skeletal muscle development and demonstrate previously unknown isoform-specific functional requirements for activation of calcineurin A transcriptional targets.

## Results

### Transcriptome changes predominate within the first two weeks of postnatal skeletal muscle development

To identify changes in gene expression and alternative splicing during postnatal skeletal muscle development, we performed RNA-seq using RNA from gastrocnemius muscle at E18.5, PN2, PN14, 28, and adult (22 weeks) including a biological replicate for PN14. Males were used for all time points except E18.5 for which both male and female animals were used. We obtained >160 million 100 bp paired end reads per sample with at least 92% of reads mapping to the mouse genome (Table 1). The PN14 biological replicates revealed strong correlations for both gene expression and alternative splicing indicating high levels of reproducibility (r^2^ = 0.99 and 0.90, respectively, Figure 1A and 1B). Splicing transitions predicted by RNA-seq were validated by RT-PCR by comparing the change in percent spliced in (ΔPSI) identified by RNA-seq and by RT-PCR between PN2 and PN28 (r^2^ = 0.80, Figure 1C-E). The overall results indicate that our RNA-seq data reflects the transcriptome changes occurring *in vivo* during postnatal skeletal muscle development.

**Table 1.** Mouse tissue samples used for RNA-seq. Samples were pooled for E18.5 and PN2 to obtain sufficient quantities of RNA.

| Sample | Tissue | Age | Sex | # of mice | Mapped reads | % mapped |
| --- | --- | --- | --- | --- | --- | --- |
| E18.5 | Gastrocnemius | E18.5 | M/F | 21 | 377,538,308 | 93.00% |
| PN2 | Gastrocnemius | PN2 | M | 6 | 381,045,805 | 92.20% |
| PN14 Replicate #1 | Gastrocnemius | PN14 | M | 1 | 321,001,586 | 92.60% |
| PN14 Replicate #2 | Gastrocnemius | PN14 | M | 1 | 379,096,597 | 93.20% |
| PN28 | Gastrocnemius | PN28 | M | 1 | 377,248,209 | 93.30% |
| Adult | Gastrocnemius | 22 weeks | M | 1 | 381,729,569 | 93.50% |

**Figure 1.**
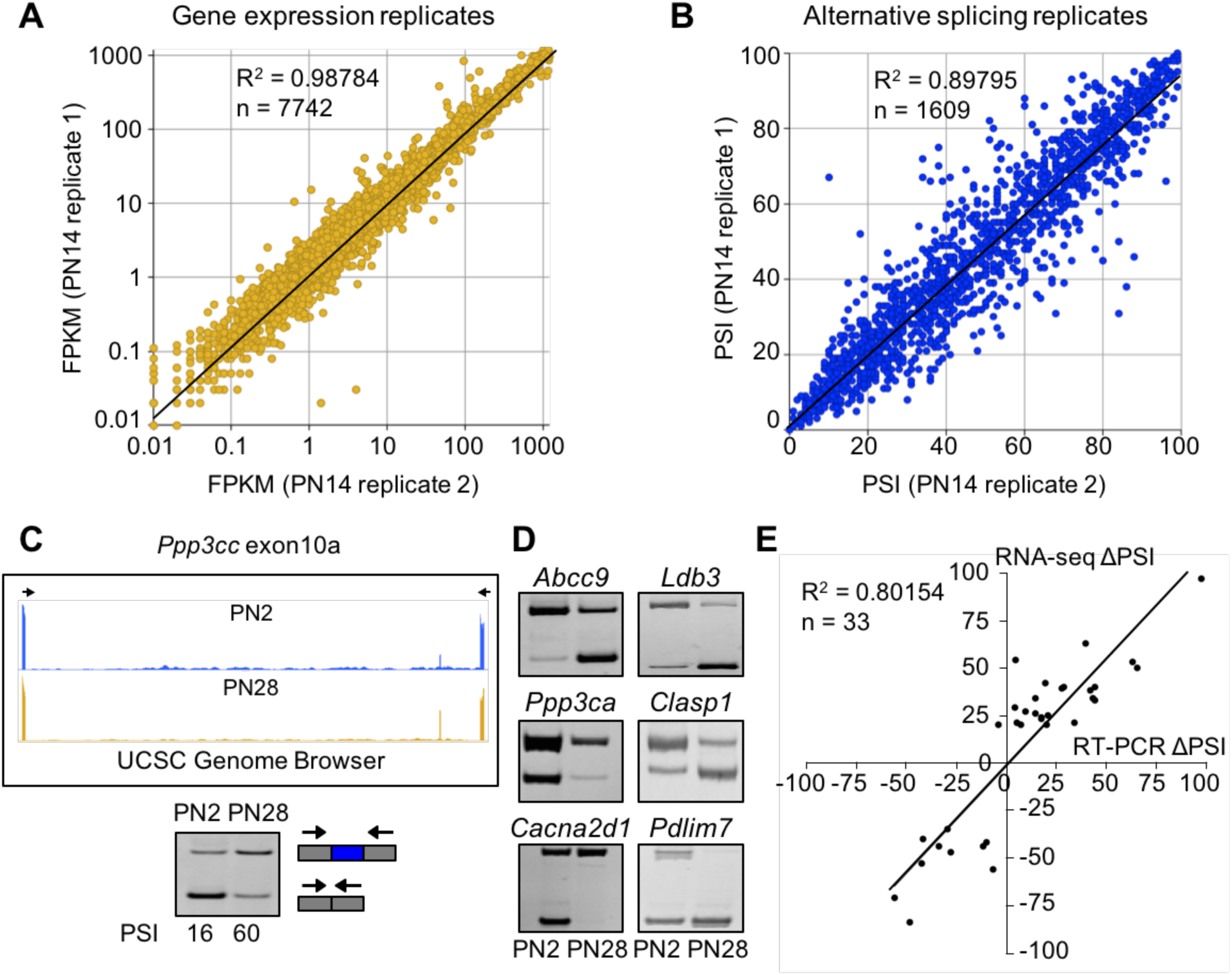
RNA-seq data is high quality and reproducible. (A and B) Biological replicates for PN14 were compared to analyze variation in gene expression (A) and alternative splicing (B) using Cufflinks and MISO, respectively. (C) RT-PCR to quantitate splicing use primers that anneal to the constitutive exons flanking an alternative exon, displayed on the UCSC Genome Browser (above) to determine the percent spliced in (PSI). (D) RT-PCR splice products for alternative splicing events comparing PN2 to PN28. (E) Plot comparing PN2 to PN28 RNA-seq ΔPSI values and ΔPSI values obtained by RT-PCR.

We identified 4417 genes showing differential expression (±2.0-fold change) and 721 events showing differential splicing (ΔPSI of ±15%) comparing E18.5 to adult skeletal muscle. For both gene expression and alternative splicing, the interval between PN2 and PN14 was the most dynamic time period with regard to the numbers of genes undergoing transitions. The analysis is affected by the differences in interval length between times points (E18.5 to PN2 vs. PN28 to adult) but even after correcting for differences in interval duration, the largest number of genes change expression between PN2 and PN14 (Figure S1A). From E18.5 to adult time points, 56% of differential gene expression changes occurred between PN2 and PN14 (3315 genes) while expression of 636 genes (11%) changed between PN14 and PN28, 481 genes (8%) changed in gene expression from E18.5 to PN2 and 1496 genes (25%) changed expression between PN28 and adult (Figure 2A).

**Figure 2.**
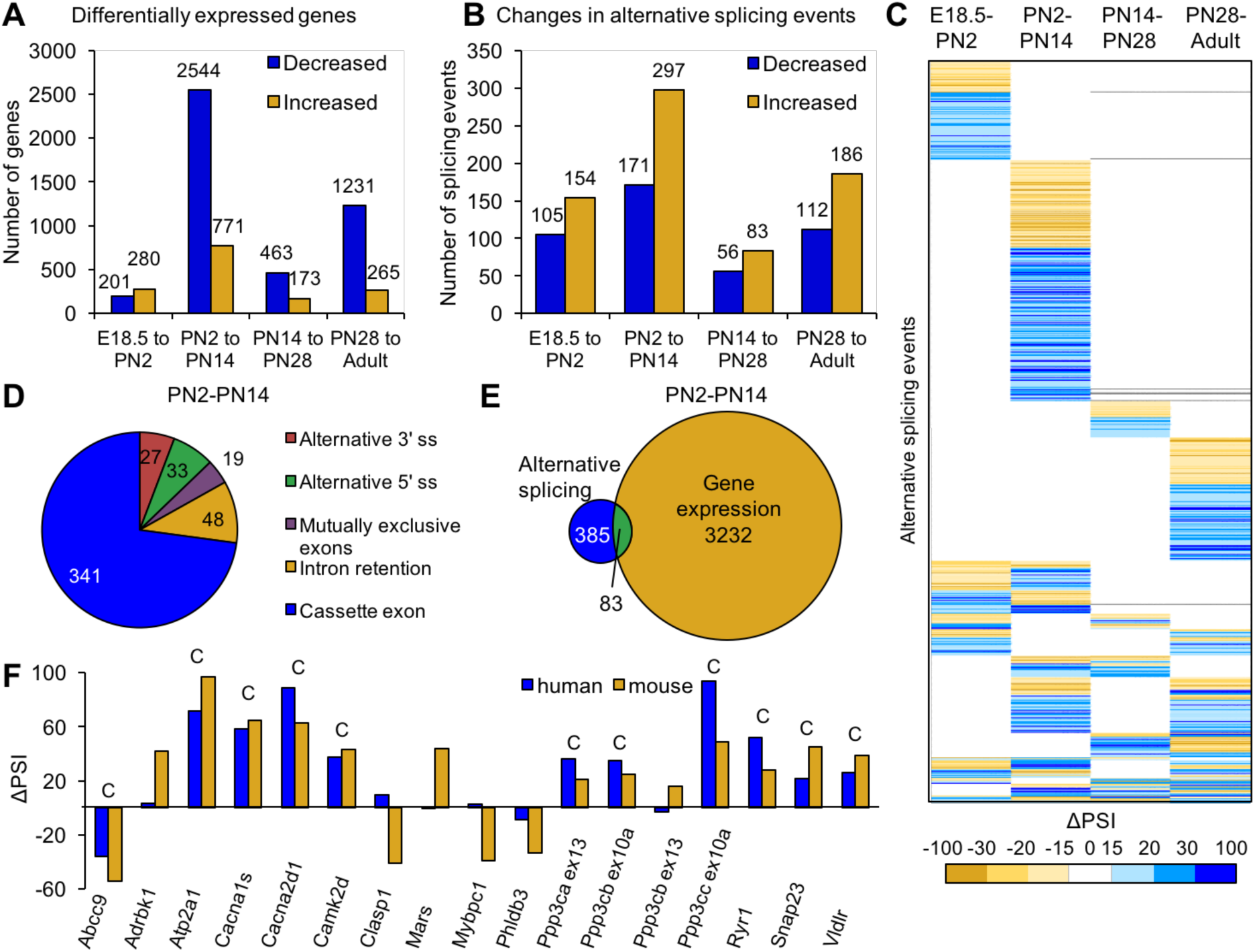
Postnatal gene expression and alternative splicing transitions in mouse skeletal muscle are largely independent, temporally restricted and conserved. (A) Genes with ≥2.0-fold increase or decrease in expression between E18.5 and adult. (B) Splicing events with ΔPSI ≥ ± 15% between E18.5 and adult. (C) Heat map of alternative splicing transitions between four time intervals. Most gene expression and splicing transitions occur between PN2 and PN14. (D) Splicing patterns of events with 15% ΔPSI or greater between PN2 and PN14. (E) Venn diagram of genes with gene expression changes (2.0-fold or greater) compared to alternative splicing transitions (15% ΔPSI or greater) between PN2 and PN14. (F) Conservation of splicing transitions during mouse and human skeletal muscle development. The ΔPSI between PN2 to PN28 mouse gastrocnemius samples were compared to the ΔPSI between gestation week 22 to adult human skeletal muscle by RT-PCR. Events showing a 15% ΔPSI or greater in the same direction in mouse and human samples were scored as conserved (indicated by C).

The numbers of proliferating satellite cells decrease during early postnatal skeletal muscle development such that a portion of the transcriptome changes are likely to reflect changes in cell population rather than transitions within established myofibers. The RNA-seq data shows that the expression of markers of activated satellite cells is relatively low even at PN2 while the changes in expression of myofiber markers (myogenin, desmin, and MyHC IIb) are robust (Figure S2). These results are consistent with the contention that the dynamic transcriptome changes reflect transitions within established myofibers with minimal contributions from a changing satellite cell population.

To identify the timing of splicing transitions, we compared ΔPSIs of the four developmental intervals (Figure 2B). Similar to differential gene expression, the interval between PN2 and PN14 has the largest number of splicing transitions (64%). Of the 768 splicing events that occur between E18.5 and adult, 32% occur specifically between PN2 and PN14 while only 13% occur between E18.5 and PN2, 5% occur specifically between PN14 and PN28, and 16% occur specifically between PN28 and adult. Sixty seven percent of the splicing events with a ΔPSI ≥ 15% undergo a transition during only one time interval (PN2-14) (Figure 2C). These results indicate that there is enrichment for alternative splicing changes within the first two weeks after birth, 77% of which are cassette exons from PN2 to PN14 (Figure 2D).

While there are large numbers of transitions for both alternative splicing and gene expression between PN2 and PN14, there is little overlap in the genes that undergo these transitions. Of the genes that undergo alternative splicing transitions between PN2 and PN14, only 18% also showed differential gene expression indicating that the majority of alternative splicing changes are within genes that do not significantly change expression (Figure 2E). Of the 768 differential splicing events that between E18.5 and adult, 31% of alternative splicing events do not exhibit differential gene expression and 58% of differential alternative splicing events are within genes that show differential expression only during a separate time interval (Figure S1B). This indicates that differential gene expression and alternative splicing are both dynamic throughout development, but are typically regulated at different times. Since the majority of splicing changes are within open reading frames (see below), for many genes there is a major impact on protein isoform transitions rather than a change in gene output.

To determine the level of conservation of validated splicing transitions, we performed RT-PCR using RNA from PN2 and PN28 mouse skeletal muscle and human skeletal muscle RNA from 22 weeks gestation and adult. Alternative splicing events were considered to be conserved if both the mouse and human had a ΔPSI of 15% or greater in the same direction (Figure 2F). Of the 17 splicing events tested, 11 (65%) underwent a transition that was conserved. All nine events involved cassette exons that inserted or removed in-frame peptides The results suggest that the different protein isoforms that result from the alternative splicing transitions have conserved physiological functions.

### In-frame alternative exons predominate in the ORF while out-of-frame alternative exons contain coding and untranslated regions

To determine the impact of alternative splicing on the protein isoforms expressed during postnatal development, we examined the distribution of cassette alternative exons within the spliced mRNAs and the effect of regulated splicing on the reading frame. For alternative exons with a ΔPSI of 15% or greater from PN2 to PN28 and in genes that contain at least three constitutive exons (223 exons), the relative position of alternative exons along the length of mRNA was determined by dividing the exon number of the alternative exon by the total number of exons then multiplying by 100 to derive the exon order relative to the mRNA 5’ end (percent from the 5’ end). Alternative exons were also separated based on whether or not they are a multiple of three nucleotides since the latter change the reading frame. Alternative exons that are a multiple of three were predominantly found to maintain the open reading frame (ORF) (129 exons of 135 considered) causing either an internal insertion or deletion of amino acids (Figure 3A and 3B). Six exons either created translation start or stop codons or altered the 5’ UTR. Alternative exons that are not a multiple of three (88 exons) were enriched near the 5’ or 3’ ends of the mRNA either exclusively within the UTRs (22 exons) or affecting the ORF to produce alternative N- or C-termini (66 exons) (Figure 3C and 3D). Therefore the majority of alternative splicing transitions produce fetal and adult protein isoforms differing by internal peptide segments. Less common, but still prevalent, are transitions that produce different N- or C-termini. The least common are alternative exons that affect untranslated regions.

**Figure 3.**
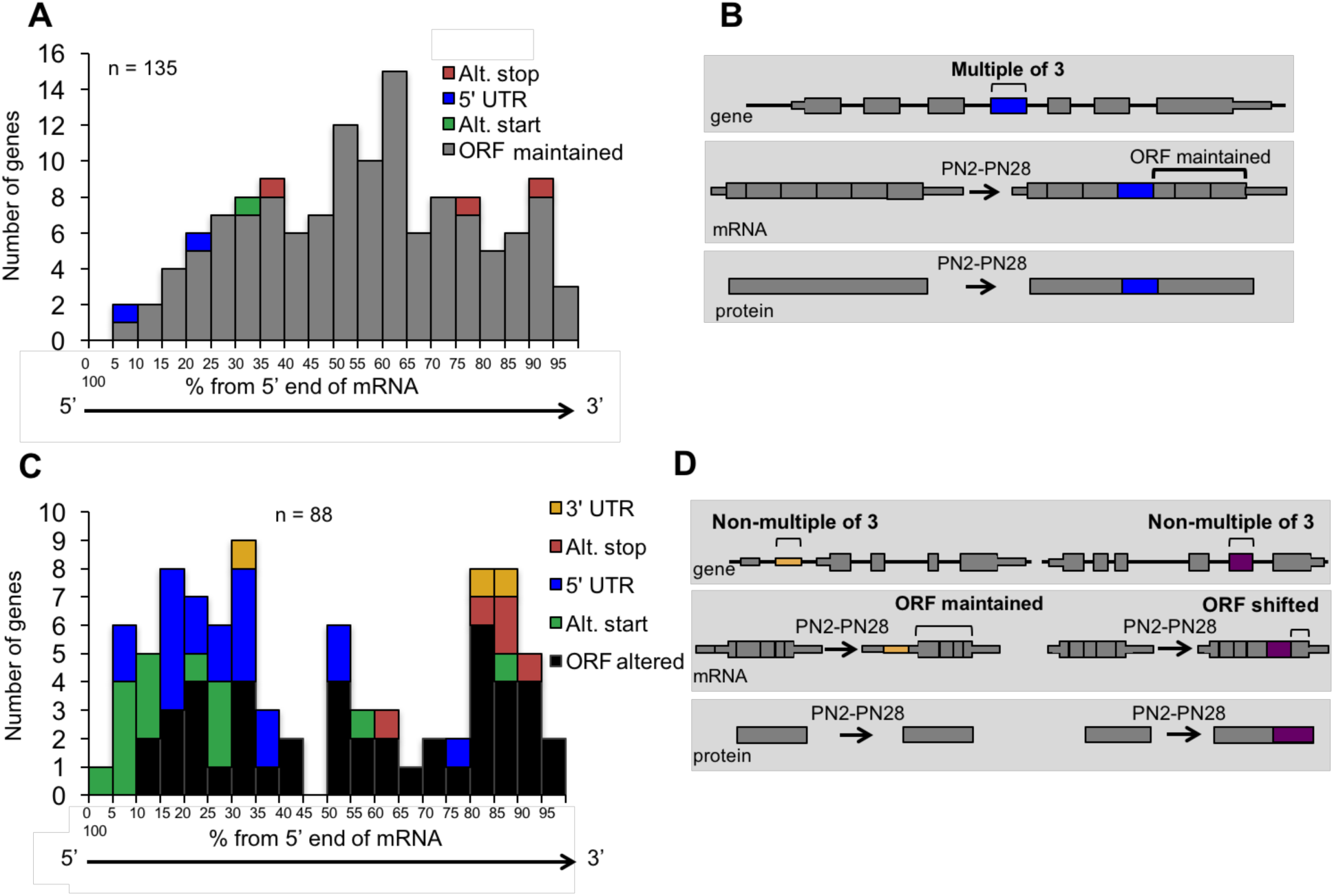
Distribution of cassette alternative exons within mRNAs. The graphs display the number of genes and the relative position of the alternative cassette exon from the 5’ end of the spliced mRNA. The relative exon position is expressed as the percent of the total number of exons. The alternative exons analyzed (223 exons) have a ΔPSI of 15% or greater from PN2 to PN28 and contain at least 3 constitutive exons. (A) In-frame alternative exons that maintain the reading frame. Colors indicate whether the alternative exon contains a translational stop codon (red), start codon (green), only 3’ UTR sequence (yellow), only 5’ UTR sequence (blue), or if the exon is within the open reading frame (gray). (B) Representation of potential protein-coding consequences of in-frame alternative exons. (C) Out of frame alternative exons that shift the reading frame. D. Representation of potential protein-coding consequences of alternative exons that are a non-multiple of three.

### Genes that undergo postnatal splicing transitions during skeletal muscle development are enriched for calcium handling functions

Ingenuity analysis of genes that undergo changes in gene expression and alternative splicing between PN2 and PN28 identified essentially non-overlapping functional categories (Figure 4). Specifically, genes that undergo differential expression were enriched for associations with mitochondrial function while alternative splicing transitions were enriched for genes associated with calcium handling, endocytosis, and cell junction categories (Figure 4A and 4B). Although calcium-related categories included 90 genes for both gene expression and alternative splicing (75 for gene expression, 21 for alternative splicing), only six genes were found to have significant expression and splicing changes. Table 2 lists the calcium handling genes that undergo ΔPSI ≥15 point change in alternative splicing between PN2 and PN28. Calcium handling is critical to striated muscle contractility and homeostasis and our results indicate that a large fraction of calcium handling genes undergo splicing transitions that affect the coding potential of these genes (Table 2 and Figure 5). These results suggest an important role for alternative splicing transitions from fetal to adult isoforms in calcium handling genes during postnatal skeletal muscle development.

**Figure 4.**
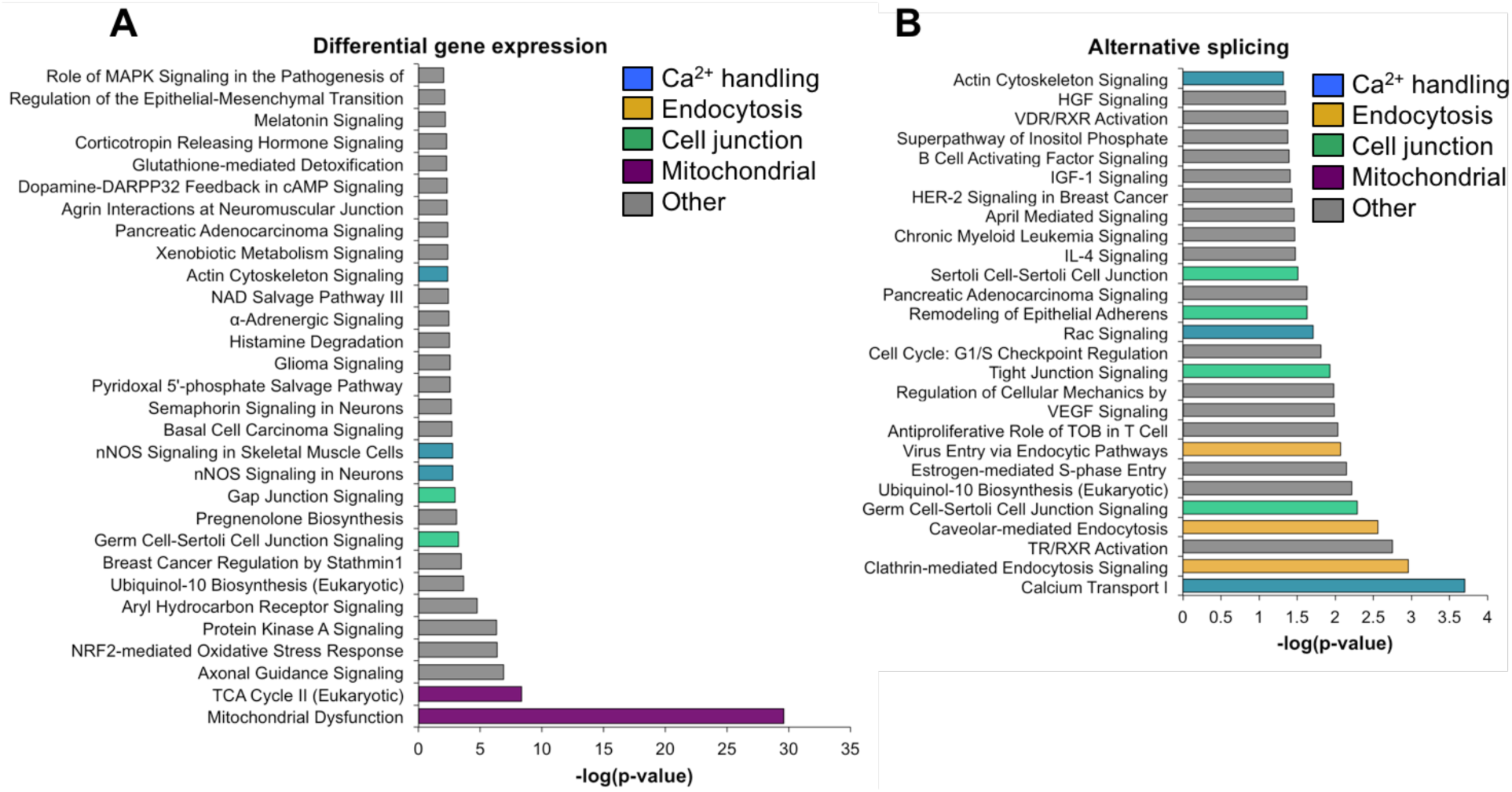
Gene ontology analysis for differential gene expression and alternative splicing. Ingenuity analysis was performed for both gene expression and alternative splicing, with a significance cut-off of −log(1.3). (A) Ingenuity analysis for gene expression differences between PN2 and PN28 (2-fold cut-off). The top 30 GO terms are displayed. For significant GO terms not shown, none of the highlighted, colored terms were present. (B) Ingenuity analysis for alternative splicing for genes with a 15% ΔPSI or greater between PN2 and PN28. All significant GO terms are shown.

**Table 2.** Calcium handling genes with alternative splicing transitions during skeletal muscle development. Listed are calcium-handling genes from the GO analysis (Figure 3). The ΔPSI from PN2 to PN28 are displayed along with the size of alternative exon, effect on the reading frame, the relative location of the exon, and predicted proteincoding consequence.

| Gene Symbol | Gene | ΔPSI | Size of exon (bp) | In frame? | Alt. exon position | Effect on coding |
| --- | --- | --- | --- | --- | --- | --- |
| Atp2a1 | ATPase, Ca<sup>++</sup> transporting, cardiac muscle, fast twitch | 97 | 42 | Y | 22 of 23 | C-term. |
| Atp2a3 | ATPase, Ca<sup>++</sup> transporting, ubiquitous | 20 | 73 | N | 21 of 22 | C-term. |
| Atp2b3 | ATPase, Ca<sup>++</sup> transporting, plasma membrane | -42 | 154 | N | 21 of 22 | C-term. |
| Cacna1s | Calcium channel, voltage-dependent, L type, alpha 1S subunit | 50 | 57 | Y | 29 of 44 | Insertion |
| Cacna2d1 | Calcium channel, voltage-dependent, alpha 2/delta subunit 1 | 53 | 57 | Y | 19 of 39 | Insertion |
| Calu | Calumenin | -27 | 194 | Y | 3 of 7 | Mutually exclusive |
| Camk2b | Calcium/calmodulin-dependent protein kinase II beta | 62 | 129 | Y | 13 of 17 | Insertion |
| Camk2d | Calcium/calmodulin-dependent protein kinase II delta | 34 | 89 | N | 19 of 20 | C-term. |
| Camsap1 | Calmodulin regulated spectrin-associated protein 1 | 25 | 33 | Y | 5 of 18 | Insertion |
| Cask | Calcium/calmodulin-dependent serine protein kinase | -44 | 69 | Y | 14 of 21 | Deletion |
| Chrne | Cholinergic receptor, nicotinic, epsilon (muscle) | -18 | 107 | N | 5 of 12 | Change of a.a. |
| Kcnn1 | Potassium intermediate/small conductance calcium-activated channel, subfamily N, member 1 | -35 | 111 | Y | 6 of 10 | Deletion |
| Mef2d | Myocyte enhancer factor 2D | -31 | 138 | Y | 4 of 12 | Mutually exclusive |
| Nfatc3 | Nuclear factor of activated T-cells, cytoplasmic, calcineurin-dependent 3 | 20 | 104 | N | 10 of 11 | C-term. |
| Ppp3ca | Protein phosphatase 3, catalytic subunit, alpha isozyme (calcineurin Aα) | 30 | 30 | Y | 13 of 14 | Insertion |
| Ppp3cb | Protein phosphatase 3, catalytic subunit, beta isozyme (calcineurin Aβ) | 25 | 27 | Y | 11 of 15 | Insertion |
| Ppp3cb | Protein phosphatase 3, catalytic subunit, beta isozyme (calcineurin Aβ) | 16 | 30 | Y | 14 of 15 | Insertion |
| Ppp3cc | Protein phosphatase 3, catalytic subunit, gamma isozyme (calcineurin Aγ) | 49 | 27 | Y | 11 of 15 | Insertion |
| Ryr1 | Ryanodine receptor 1 (skeletal) | 39 | 18 | Y | 83 of 106 | Insertion |
| Tnnt3 | Troponin T type 3 (skeletal, fast) | 22 | 41 | Y | 16 of 17 | Mutually exclusive |
| Trdn | Triadin | 18 | 60 | Y | 9 of 36 | Insertion |

**Figure 5.**
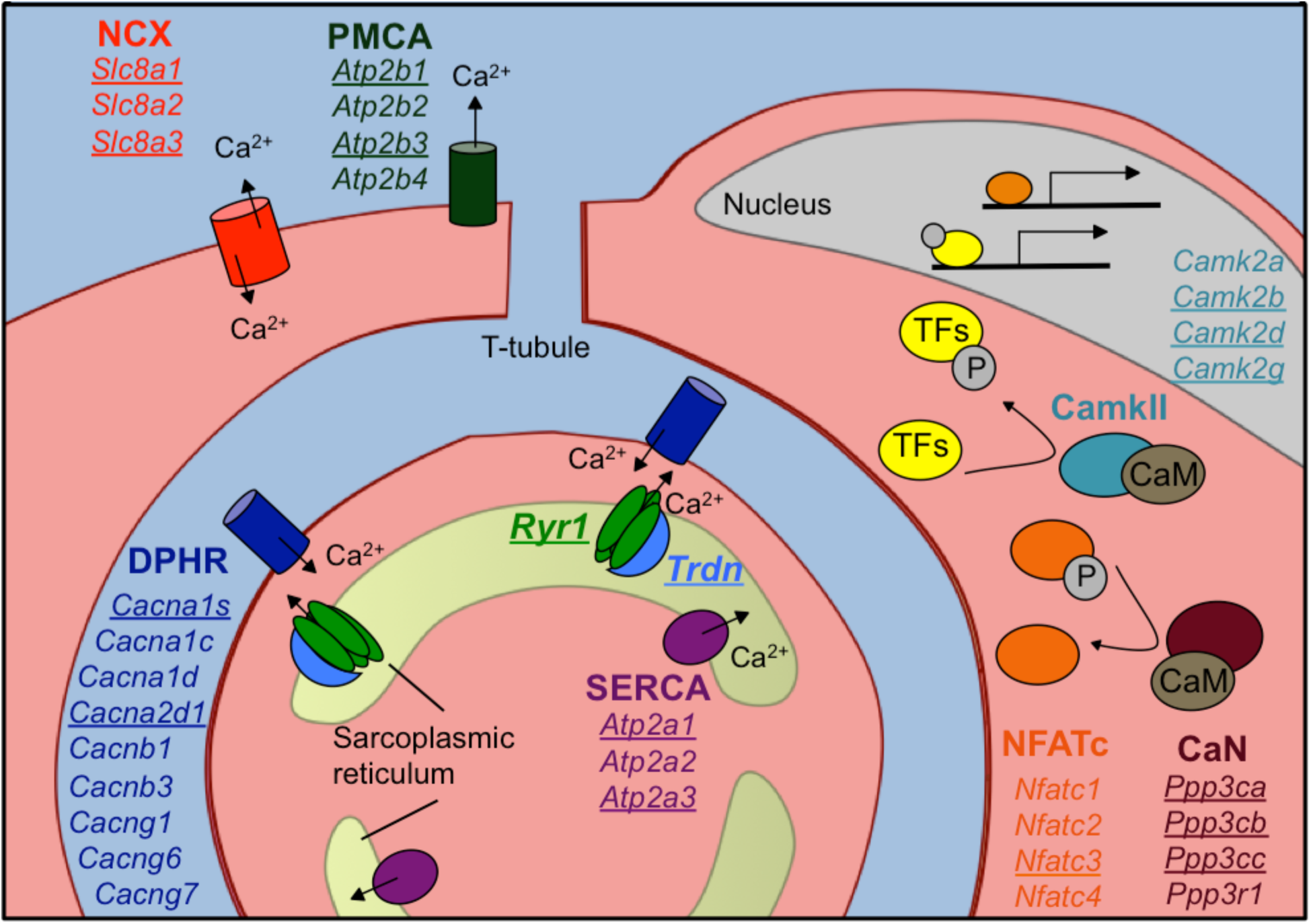
Calcium handling genes that undergo alternative splicing transitions in postnatal skeletal muscle development. Diagram of calcium handling genes that are expressed in skeletal muscle. Genes with 15% or greater ΔPSI from PN2 to PN28 are underlined and include members of several calcium channels: NCX (sodium calcium exchanger), PMCA (plasma membrane Ca^2+^-ATPase), SERCA (sarco/endoplasmic reticulum Ca^2^+-ATPase), and RYR (ryanodine receptor); triadin (Trdn) which associates with Ryr1, junctin (Asph) and FKBP12 (Fkbp1a). Signaling cascades that are affected by alternative splicing include Ca^2^+/camodulin (CaM)-dependent calcineurin (CaN) and calmodulin-dependent protein kinase II (CamkII) along with the downstream transcription factor NFATc. NFATc and transcription factors (TFs) regulated by CamkII activate genes for hypertrophy and fiber type specification.

### CELF and MBNL proteins regulate distinct sets of calcium handling genes

CELF and MBNL proteins are involved in alternative transitions during normal developmental and in skeletal muscle disease (13,14,32,36,37). Of the six Celf paralogs and three Mbnl paralogs in mice, Celf1, Celf2, Mbnl1, and Mbnl2 are expressed in postnatal and adult skeletal muscle and therefore comprise the totality of CELF and MBNL activities during postnatal development. Western blot analysis of protein expression during postnatal development of gastrocnemius muscle demonstrated that Celf1, Celf2, and Mbnl2 protein levels decrease dramatically between PN7 and PN14 (Figure 6A). Mbnl1 protein expression is reduced after PN7 and published results indicate that Mbnl1 undergoes translocation to the nucleus during postnatal skeletal muscle development (32). These results indicate a strong correlation between the most dynamic period of splicing change and differential protein expression of these two families of splicing regulators.

**Figure 6.**
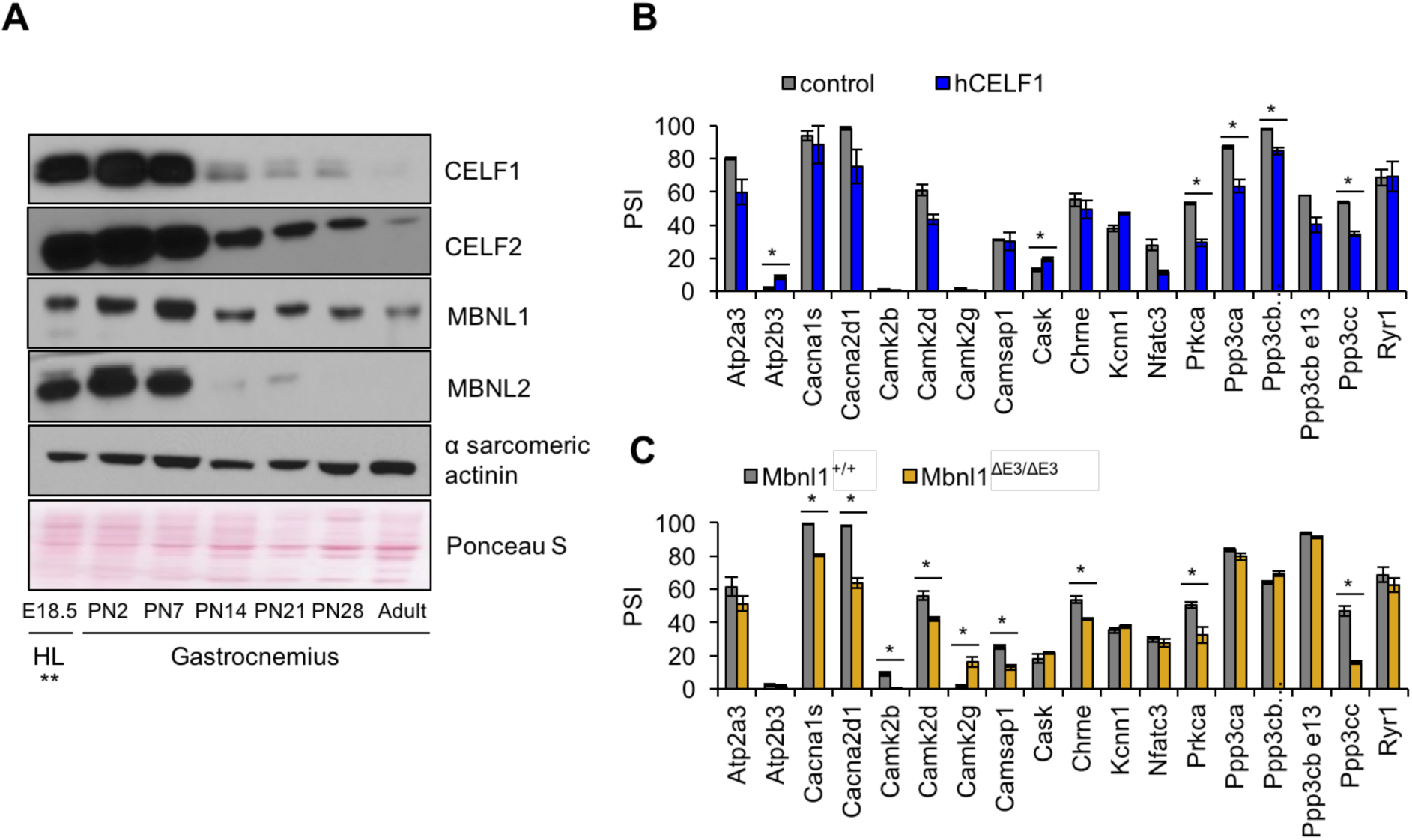
Postnatal down regulation of CELF and MBNL alternative splicing regulators promote splicing transitions of calcium handling genes. (A) Western blot of Celf1, Celf2, Mbnl1, and Mbnl2 during gastrocnemius postnatal development. **All E18.5 samples are from hindlimb (HL) except Celf1 blot which is E18.5 gastrocnemius. Ponceau S and α sarcomeric actinin serve as loading markers. (B) Comparing PSI of control mice (MDAFrtTA +dox) and human CELF1 overexpressing mice (MDAFrtTA/TRECUGBP1 +dox) (C57BL6/DBA;FVB). (C) Comparing PSI of wild type and Mbnl1 KO mice, Mbnl1^ΔE3/ΔE3^ (FVB). Single asterisk (*) denotes P<0.05 using student T-test, n = 3 mice per group.

To determine whether changes in Celf1 and Mbnl1 protein levels affect postnatally regulated alternative exons in calcium handling genes, we analyzed splicing of these genes in skeletal muscle from transgenic mice induced to overexpress human CELF1 in adult animals (MDAFrtTA/TRECUGBP1 +dox vs. control MDAFrtTA +dox) and from adult *Mbnl1^ΔE3/ΔE3^* knock out mice (*Mbnl1^ΔE3/ΔE3^* vs. control *Mbnl1^+/+^*) (Figure 6B-C and Figure S3) (38,39). Overexpression of CELF1 and loss of Mbnl1 produced six and nine significant splicing changes, respectively, (15 genes affected of 18 tested) all of which reversed the direction observed during postnatal development. Only two genes, *Ppp3ca* and *Ppp3cc* responded to both gain of CELF1 and loss of Mbnl1. These results indicate that CELF and MBNL proteins are important contributors to regulated splicing within genes involved in calcium handling during postnatal skeletal muscle development.

### All three calcineurin A genes undergo fetal to adult protein isoform transitions during postnatal development

To determine the functional consequences of postnatal splicing transitions, we focused on the calcineurin A genes (*Ppp3ca, Ppp3cb*, and *Ppp3c*). Calcineurin is a calcium sensitive phosphatase affecting fiber type in skeletal muscle by dephosphorylating Nuclear Factor of Activated T-cells component (Nfatc) proteins leading to its nuclear translocation and Nfatc-mediated transcriptional activation of a select subset of genes. It is a heterodimer containing a catalytic subunit, calcineurin A, and regulatory subunit, calcineurin B. All three calcineurin A paralogs expressed in skeletal muscle show postnatal splicing transitions with a ΔPSI of at least 15% (Table 2, Figure 5, and Figure 7A) while the two calcineurin B genes do not (data not shown). The functional consequences for the calcineurin A splicing events have not been characterized.

**Figure 7.**
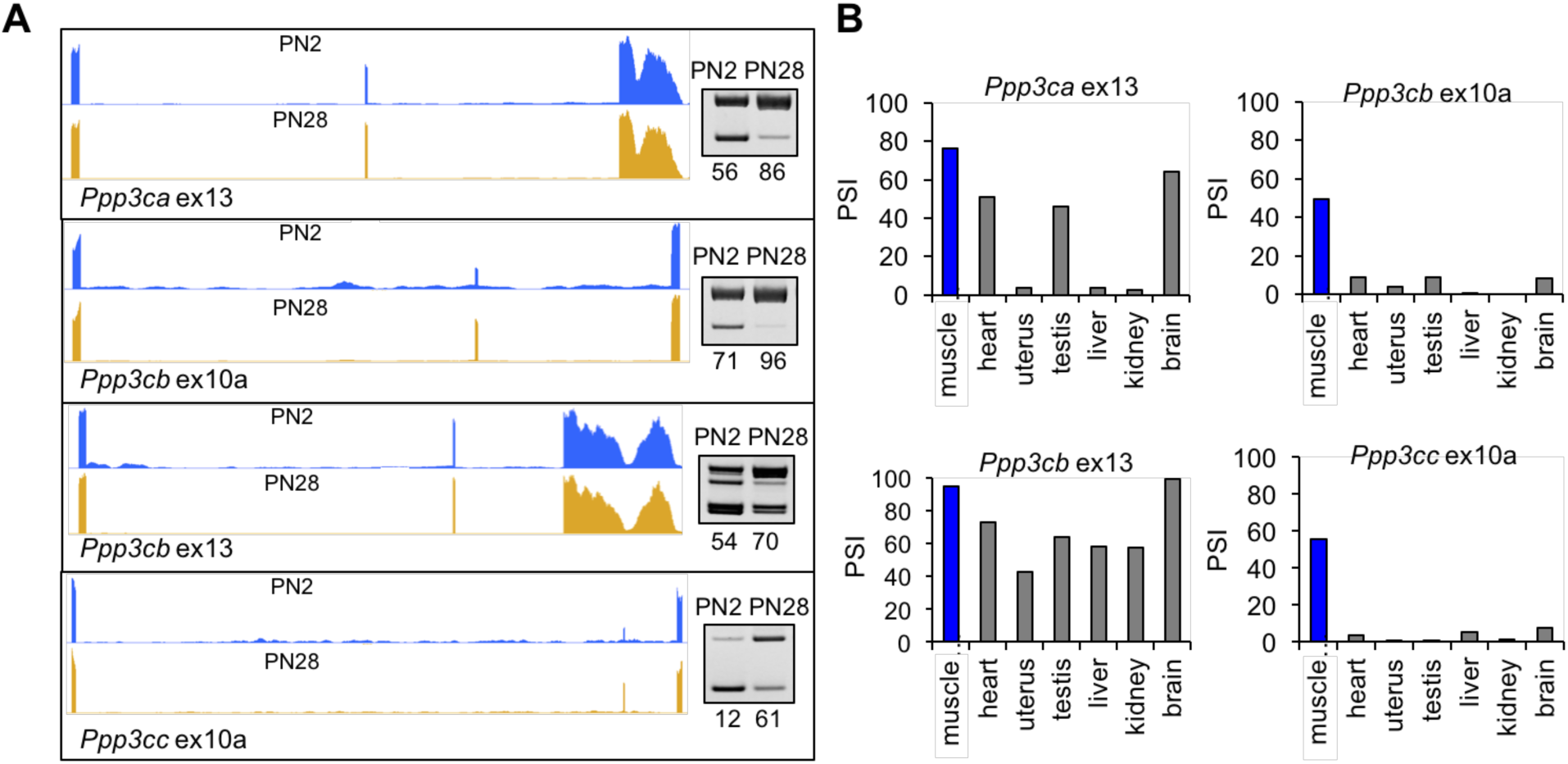
Calcineurin A splicing during PN development in different tissues. (A) UCSC Genome Browser displays of calcineurin A alternative exons (*Ppp3ca* ex13, *Ppp3cb* ex10a, *Ppp3cb* ex13, *and Ppp3cc* ex10a) side-by-side with RT-PCR of RNA from PN2 and PN28. (B) RT-PCR comparing inclusion of calcineurin A alternative exons in diverse adult tissues.

*Ppp3cb* and *Ppp3cc* each have an homologous 27 base pair alternative exon 10a with in-frame insertions encoding peptide segments with low sequence identity (Figure S4A). *Ppp3ca* does not have a protein sequence equivalent to exon 10a. In contrast, *Ppp3ca* and *Ppp3cb* each have an alternatively spliced 30 base pair exons 13 encoding nearly identical amino acid sequences (Figure S4A). Exon 13 for *Ppp3cc* is constitutively included and encodes a peptide sequence that is 40% and 50% identical to *Ppp3ca* and *Ppp3cb*, respectively (Figure S4A). All four alternative exons have increased inclusion during postnatal development. In addition, exons 10a of *Ppp3cb* and *Ppp3cc* are included specifically in adult skeletal muscle while exons 13 of *Ppp3ca* and *Ppp3cb* are included in other adult tissues including heart, testis, and brain (Figure 7B).

Calcineurin A contains a calcineurin B binding domain, calmodulin binding domain, phosphatase domain, and an auto-inhibitory domain (40). Alternative exons 10a of *Ppp3cb* and *Ppp3cc* are adjacent to and upstream of the calmodulin-binding domain, and exons 13 of *Ppp3ca* and *Ppp3cb* are adjacent to and upstream of the autoinhibitory domain (Figure S4B). These positions suggest a potential effect on the functions of these domains. In addition, the conservation of the postnatal transitions between mouse and human suggests that the isoform transition is functionally relevant to tissue remodeling during the fetal to adult transition (Figure 2F). Inclusion of calcineurin A alternative exons during postnatal development is not strongly affected by fiber type since the splicing transitions are similar between gastrocnemius and soleus muscles, which have different proportions of fast and slow fiber muscles (Figure S4C).

### Re-directed calcineurin A splicing reveals an isoform-specific effect on Nfatc2 and Nfatc3 activation

To determine the specific effects of calcineurin A redirected splicing on downstream signaling, we used the mouse C2C12 myogenic cell line. C2C12 cells transition from proliferative myoblasts to fused myotubes upon withdrawal of growth factors. Differentiating C2C12 cultures reproduce the developmental inclusion of exons 13 of *Ppp3cb* and *Ppp3ca* but not *Ppp3cc* or inclusion of exon 10a of *Ppp3cb*. We used morpholino antisense oligonucleotides (ASO) to redirect splicing of exons 13 of *Ppp3cb* and *Ppp3ca* through C2C12 differentiation (Figure 8A). Immunofluorescence staining was then used to determine the effects on Nfatc family members that are expressed in differentiated myotubes. Nfatc2 and particularly Nfatc3 showed significantly increased nuclear localization in myotubes with calcineurin A redirected splicing while Nfatc1 localization was unchanged (Figure 8B-C, Figure S5A). The Nfatc3 immunofluorescence signal was validated by knockdown in C2C12 myotubes (Figure S5B). These results demonstrate that expression of the endogenous fetal isoforms of *Ppp3ca* and *Ppp3cc* is sufficient to activate translocation of two of the three Nfatc proteins, Nfatc3.

**Figure 8.**
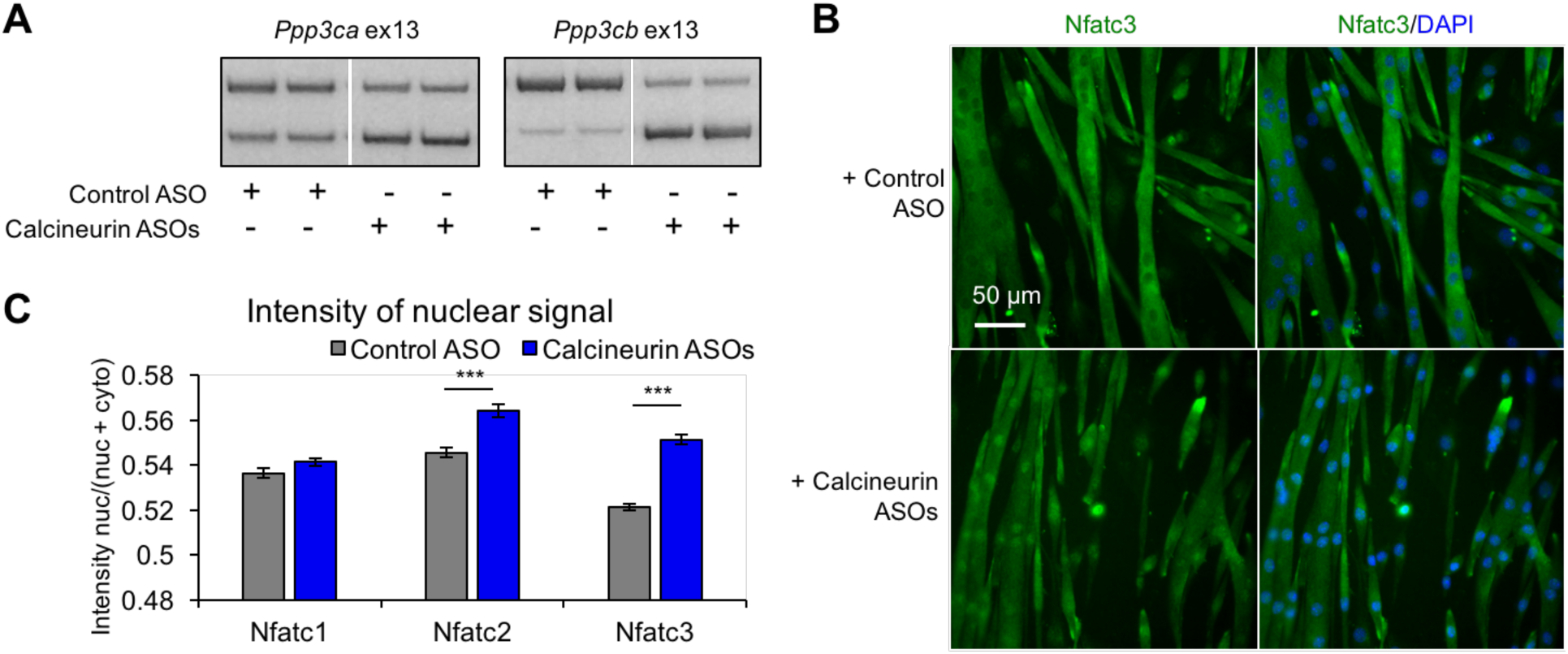
Redirected calcineurin A splicing in differentiated C2C12 myotubes. (A) RT-PCR of redirected splicing of calcineurin A events in differentiated C2C12 cells. Two of the four calcineurin A splicing events are present in C2C12. For calcineurin A ASO, 9 μΜ Ppp3ca ex13 3’ss and 15 μΜ Ppp3cb ex13 5’ss morphlinos were delivered, and for control ASO, 24 μΜ standard control morpholino were delivered. (B) Immunofluorescence of Nfatc3 in differentiated C2C12 after morpholino delivery. (C) Quantification of nuclear immunofluorescent signal of Nfatc1, Nfatc2, and Nfatc3. The nuclear intensity was determined by (nuclear intensity)/(nuclear intensity + cytoplasmic intensity). Three asterisks (***) denotes P<0.001 significance by one-way ANOVA analysis, n = ∼500 nuclei per condition.

To determine the physiological impact of the calcineurin A splicing transitions in adult skeletal muscle, we used ASOs delivered into the flexor digitorum brevis (FDB) foot pad muscle to redirect all four calcineurin A exons to the fetal splicing pattern of exon skipping. This approach allows testing the functions of specific endogenous protein isoforms without changing the overall expression level. We found that the efficiency of redirected splicing remains high for at least four weeks and three weeks provides sufficient time for the muscle to recover from the ASO delivery procedure (29).Three weeks after delivery of redirecting ASOs or non-targeting control ASOs, redirected splicing was assayed by RT-PCR and all four exons were found to have undergone a nearly complete switch to the fetal pattern while non-targeting ASOs had no effect (Figure 9A). Four non-targeted developmental splicing events were assayed (*Atp2a1, Cacna1s, Prkca, Ppp1r12b*) and none showed significant differences following ASO injection, demonstrating the absence of non-specific effects (Figure 9B).

**Figure 9.**
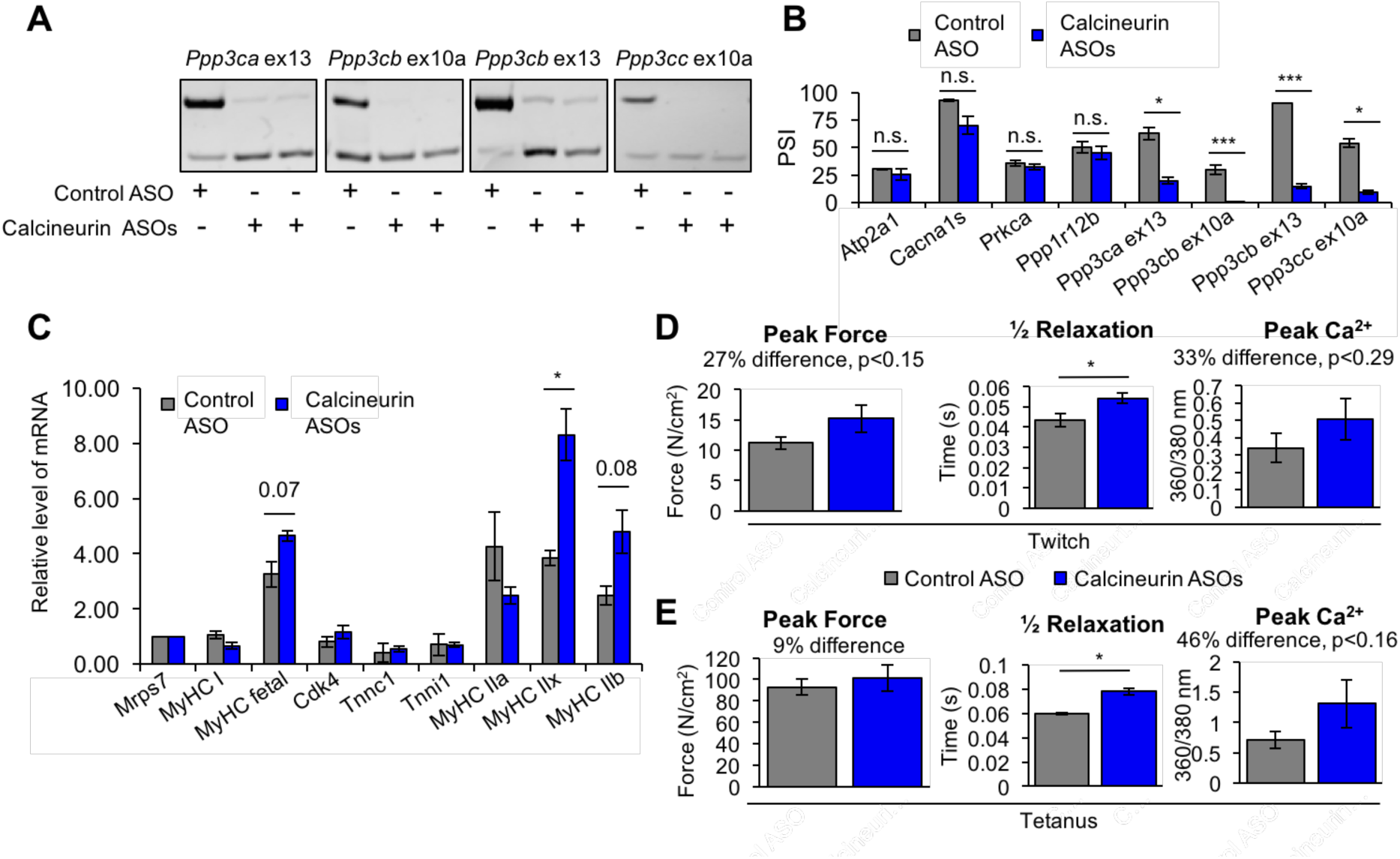
Redirected calcineurin A splicing in adult FDB muscle. (A) Confirmation of redirected splicing in adult FDB muscle by RT-PCR. ASO morpholinos were administered to the FDB muscle (Figure S7). (B) RT-PCR of targeted (calcineurin A) and control (Atp2a1, Cacna1s, Prkca, and Ppp1r12b) splicing events. Single asterisk (*) denotes P<0.05, *** denotes P<0.001, and n.s. denotes no statistical difference by student T-test, n = 4 mice per group. (C) Relative mRNA levels of mRNAs from Nfatc target genes in FDB muscle by RT-PCR. Single asterisk (*) denotes P<0.05 and marked are transcriptional targets nearing significance by student T-test, n = 3 mice per group. (D) Force and calcium analysis after twitch stimulus. Peak force and half relaxation time were measured for force and peak calcium for calcium. Single asterisk (*) denotes P<0.05 significance by student T-test, n = 3-4 mice per group. (E) Force and calcium analysis after tetanus stimuli. Peak force and half relaxation time were measured for force and peak calcium for calcium. Single asterisk (*) denotes P<0.05 significance by student T-test, n = 3-4 mice per group.

To assess physiological changes from redirected calcineurin A splicing, we measured parameters of force generation and calcium handling in FDB muscle bundles three weeks after ASO delivery. Redirected splicing resulted in a significant prolongation of twitch half relaxation time and a strong trend toward increased peak twitch force and peak calcium, although these changes did not reach statistical significance (Fig. 9C). There was no difference in twitch time to peak force or calcium for a twitch (Fig. S6). Following peak tetanic stimulation (150 Hz), we also found an increase, albeit smaller, increase in tetanic peak force, a rend toward increased peak calcium, and a significant prolongation of the half relaxation time (Fig 9D).

To determine whether expression of the fetal calcineurin A isoforms in adult FDB muscle affected Nfatc transcriptional activity, we assayed expression of known Nfatc-targeted genes. MyHC IIx showed a significant change while MyHC IIb, and MyHC fetal genes showed a trend that did not reach significance (Figure 9E). It is interesting to note that increased calcineurin activity typically leads to a slow fiber type gene program, and Nfatc3 knockout equally affects fast and slow MyHC transcription, yet manipulation of calcineurin A splicing promoted fast fiber type gene expression. Our results strongly suggest that the fetal isoforms of at least *Ppp3ca* and *Ppp3cb* have intrinsically higher phosphatase activity than the adult isoforms. These results also identify Nfatc3, and partially Nfatc2, as the family member that is primarily responsive to the fetal isoforms. This leads to Nfatc2 and Nfatc3 transcriptional changes producing a prolonged relaxation time of the skeletal muscle.

## Discussion

Our analysis of postnatal skeletal muscle development indicates that both transcriptional and post-transcriptional changes, particularly alternative splicing, are highly dynamic and largely specific to the first two weeks after birth. Seventy-nine percent of genes that undergo postnatal splicing transitions do not show significant changes in gene expression indicating a separation of regulated splicing and the regulation of mRNA steady state levels. That different gene sets are regulated at the level of splicing and mRNA levels has been found during other periods of physiological change including T cell activation, muscle differentiation and heart development (13,41-43). These results support a mechanism of regulation that involves transitions of fetal to adult protein isoforms rather than a change in total gene output for a relatively large subset of genes. Genes regulated by differential expression or alternative splicing were enriched for different functional categories: mitochondrial function for differentially expressed genes and calcium handling, endocytosis, and cell junctions for genes that undergo alternative splicing transitions. The results for alternative splicing during postnatal skeletal muscle development are similar to postnatal heart development in which vesicular trafficking genes were enriched among those regulated by alternative splicing (13,14).

Calcium regulates signal transduction, muscle contraction, and cellular homeostasis. Our results indicate that calcium channels, calcium-dependent phosphatases, and calcium-dependent kinases are alternatively spliced soon after birth. Several calcium handling genes have been established as alternatively spliced such as Serca1 (*Atp2a1*) and the ryanodine receptor 1 (*Ryr1*) while most, such as calcineurin A, have not been characterized (44-46). We show that all three calcineurin A genes (Ppp3ca, Ppp3cb and Ppp3cc) undergo fetal to adult protein isoform transitions within the first two weeks after birth. Redirect splicing to the fetal patterns demonstrate that the fetal protein isoforms do not fulfill the functional requirements of adult skeletal muscle tissue in that both a significant increase in time to relaxation and a physiologically sizable, although not statistically significant, increase in peak force and calcium. We propose that the mechanisms of altered adult muscle function are due to altered regulation of Nfatc proteins by calcineurin A. We show that redirected splicing of Ppp3ca and Ppp3cb in differentiated C2C12 myotubes leads to nuclear localization of Nfatc3 and Nfatc2, two of the three Nfatc proteins expressed in these cultures. Our results are consistent with previous results showing that calcineurin overexpression in C2C12 causes nuclear localization of Nfatc3 (47). Overall our results support a model in which the fetal calcineurin isoforms have higher intrinsic phosphatase activity on Nfatc3 and Nfatc2 thereby promoting its activation. Consistent with this model, we found that at least one Nfatc transcriptional target was up regulated in adult muscle expressing the fetal calcineurin isoforms due to redirected splicing. Alternative splicing of calcineurin A could be a mechanism by which calcineurin dephosphorylates specifically Nfatc3 and Nfatc2 and redirects expression of a subset of gene targets not regulated by Nfatc1.

Protein segments encoded by conserved tissue specific alternative exons are enriched for disordered domains more likely to affect protein-protein interactions and contain post-translation modification (PTM) sites (20,48). Affecting protein-protein interactions or PTMs could be a means by which calcineurin splicing events affect calcineurin activity or stability. The alternative exons could affect the ability of calcineurin A to be regulated by calmodulin or through auto-inhibitory inhibition. A splice variant, calcineurin Aβ1, has a C-terminal truncation so that the auto-inhibitory domain is missing. Overexpression of this splice variant improved cardiac function in mice (49,50). Calcineurin Aβ1 was not abundant in our RNA-seq analysis of skeletal muscle development, but it does give insight to functional consequences of an alternative isoform.

We also investigated the mechanisms that regulate a subset of the dramatic splicing transitions during postnatal development. We identified abrupt changes in CELF and MBNL protein levels during postnatal skeletal muscle development that temporally correlate with the splicing transitions that occur by PN14. We used CELF1 overexpressing and Mbnl1 knockout mice to show that 72% of the calcium handling genes tested that exhibit alternative splicing transitions respond to these proteins and are likely regulated by the natural transitions in CELF and MBNL activity that occur during postnatal development. Further investigation into Celf1^-/-^ mice showed transcriptome changes, including a network of Celf1-dependent splicing, that cause heart defects in early postnatal development (51). Celf1 is also likely to be important for muscle regeneration as Celf1 protein increases after muscle injury and Celf1 splicing targets revert to fetal splicing pattern (52). Mbnl proteins negatively regulate embryonic stem cell-like pattern of alternative splicing, and overexpression of Mbnl1 in embryonic stem cells produces a differentiation-like cell alternative splicing pattern (53). Since Mbnl1 and Mbnl2 protein levels decrease postnatally, this effect on splicing of differentiated and embryonic stem cells is likely reflective of Mbnl cellular localization switch during development and not the overall protein levels of Mbnl1 or Mbnl2 (32).

A growing number of global RNA-seq analyses are revealing the extent to which conserved alternative splicing transitions are critical for tissue function (13,42,48,54,55). In this work we identify the dynamic transcriptome changes during skeletal muscle postnatal development and demonstrate the functional significance for the calcineurin A splicing transitions. The results demonstrate the importance of understanding the functional differences between fetal and adult protein isoforms and the contribution of these proteins to tissue remodeling to adult function.

## Acknowledgements

We thank Jimena Giudice (UNC) for her advice on RNA-seq analysis and FDB electroporation, Joshua Sharpe (BCM) for help on comparing alternative splicing between human and mice by RT-PCR, Kathleen Manning (BCM) for help with RNA-seq analysis, and Charles Thornton for *Mbnl1^ΔE3/ΔE3^ mice*. This project has been supported by NIH grants R01AR045653, R01HL045565, and R01AR060733 (to T.A.C.), NIH R01HG007538 and R01CA193466 (to (WL) NIH grant R01AR061370 (to G.G.R.), NIH grant T32 HL007676 (JAL), and the Muscular Dystrophy Association (T.A.C.). This project was supported by the Genomic and RNA Profiling Core at BCM and the expert assistance of the core director, Dr. Lisa D. White, Ph.D. The Integrated Microscopy Core at BCM also supported this project with funding from NIH (NCI-CA125123, NIDDK-56338-13/15), Cancer Prevention Research Institute of Texas (RP150578), and John S. Dunn Gulf Coast Consortium for Chemical Genomics.

## Materials and Methods

### Animals

Skeletal muscle tissues were isolated from FVB wild type, MDAFrtTA/TRECUGBP1, and Mbnl1^ΔE3/ΔE3^ mice. We followed NIH guidelines for use and care of laboratory animals approved by Baylor College of Medicine Institutional Animal Care and Use Committee.

### Skeletal muscle isolation and RNA extraction

Animals were anaesthetized, either decapitated (neonatal) or cervical dislocated (older than PN10), and gastrocnemius muscles were removed. Sex determination for animals PN7 and younger was confirmed by PCR using primers to Actin and Sry genes (sequences in Fig. S7). Tissue samples were flash frozen with liquid nitrogen. Total RNA was prepared using the RNeasy fibrous tissue mini kit (Qiagen).

### RNA-seq

Illumina TruSeq protocols were used to prepare libraries using total RNA (2 ug) from gastrocnemius of E18.5, PN2, PN14, PN28, and 22 week (adult) animals. The cDNA was created using the fragmented 3’-poly(A)-selected portion of total RNA and random primers. To generate the libraries, the blunt ended fragments of cDNA were attached to adenosine to the 3’-end and ligated with unique adapters to the ends. The ligated products were amplified by PCR for 15 cycles. Libraries were quantified and fragment size assessed by the NanoDrop spectrophotometer and Agilent Bioanalyzer, respectively. The libraries were amplified by qPCR to determine the concentration of adapter-ligated fragments using a Bio-Rad iCycler iQ Real-Time PCR Detection System and a KAPA Library Quant Kit. The library (11 pM) was loaded onto a flow cell and amplified by bridge amplification using Illumina cBot equipment. On a HiSeq Sequencing system, a paired-end 100 cycle run was used to sequence the flow cell.

### Computational processing and bioinformatics of RNA-seq data

For the RNA-seq alignment, paired-end RNA-seq reads were aligned to the mouse genome (mm9) using TopHat 2.0.5 (56). For the differential gene expression analysis, RSEM was used to count the number of fragments mapped into RefSeq gene models, and edgeR was used to call differentially expressed genes with a false discovery rate less than 0.05 (RSEM: accurate transcript quantification from RNA-Seq data with or without a reference genome) (43,57). The gene expression was quantified by FPKM. For the differential alternative splicing analysis, isoform levels (PSI) and Bayes factors (BI) were measured by MISO (58) with ΔPSI >= 0.1 and BI >= 10. For both differential gene expression and alternative splicing analysis, Data were analyzed through the use of QIAGEN’s Ingenuity^®^ Pathway Analysis (IPA^®^, QIAGEN Redwood City, www.qiagen.com/ingenuity).

### AS validations & human conservation by RT-PCR

Human skeletal muscle RNA samples were obtained for fetal 22-week (BioChain [R1244171-50]) and adult (Clontech [636534]). Various adult mouse tissue total RNA samples (muscle, heart, uterus, testis, liver, kidney, and brain) were obtain from BioChain. For mouse and human RNAs, 2.5 ug of RNA was used for reverse transcription (RT). RT was performed by High Capacity cDNA RT Kit (Applied Biosystem) followed by PCR (GoTaq DNA Polymerase, Promega). RT-PCR products were separated by 6% PAGE. PCR reactions involved: 95^°^C for 3 min, 25-30 cycles of 95^°^C for 45 s, 55^°^C for 45 s, 72^°^C for 45 s, and 72^°^C for 5 min. RNA-seq data was used identify alternatively splicing regions and primers (Sigma) that anneal to the constitutive flanking exons were designed. Primers are listed in Supplementary Table 6. Ethidium bromide-stained RT-PCR bands were quantified by Kodak Gel logic 2000 and Carestream Software. PSI values were calculated by densitometry using the equation: PSI = 100 × [Inclusion band/(Inclusion band + Skipping band)].

### Western blotting

FVB wild type gastrocnemius tissues were lysed in HEPES-sucrose buffer (10 mM HEPES pH 7.4, 0.32 M sucrose, 1 mM EDTA and protease inhibitors) using Bullet blender (Next Advance) and SDS (final concentration of 1%) was added before sonication (3 min at 75 V for 30 s on and 30 s off). The samples were centrifuged for 10 min at 12,000 rpm at 4^°^C. Supernatants were transferred to new tubes, and samples were diluted in loading buffer (100 mM Tris-HCl pH 6.8, 4% SDS, 0.2% bromophenol blue, 20% glycerol, 200 mM β-mercaptoethanol) then boiled for 3 min. Pierce Compatable BCA protein assay kit (Thermo Scientific) was used to quantify protein concentration after the addition of loading buffer. For each sample 40 μg of protein was loaded into a 10% SDS-PAGE gel. Proteins were transferred to membranes and blocked with 5% milk/0.1% Tween-PBS buffer for 1 hr, washed, and incubated overnight at 4^°^C with 5% milk/Tween-PBS buffer diluted primary antibodies: mouse monoclonal anti-CUGBP1 clone 3B1 (1:1000), Celf2 (Santa Cruz Biotechnology [sc-47731]-1:1000), Mbnl1 (LifeSpan Biosciences [LS-C30810]-1:1000), Mbnl2 (Santa Cruz Biotechnology [sc-136167]-1:1000), rabbit polyclonal anti-sarcomeric α-actinin (Abcam #ab72592-1:2,000). CUGBP1 and Celf2 were conjugated to HRP using Abnova Peroxidase Labeling Kit – NH_2_. Non-conjugated primary antibodies were incubated the following day for 1 h at room temperature with secondary antibodies: goat anti-mouse IgG light chain-specific HRP conjugated (Jackson Immunoresearch [#115-035-174]-1:10000) and goat anti-rabbit IgG HRP conjugated(Invitrogen, [#621234]-1:5000). Super Signal West Pico Chemilumiescent Substrate kit (Thermo Scientific) was used for developing.

### ASO injection *in vivo*

Animal protocols were approved by IACUC at Baylor College of Medicine. FVB wild type adult mice were anesthetized by isoflurane in a chamber and moved to a nose cone for injections. First, the FDB muscle was pretreated with hyaluronidase (0.5 mg/ml, 10 μl) injected subcutaneously. After 2 hours, morpholino ASOs (80 μg Ppp3ca ex13 3’ss, 20 μg Ppp3cb ex10a 3’ss, 20 μg Ppp3cb ex13 5’ss, and 80 μg Ppp3cc ex 10a 3’ss or 200 μg of standard control, 15 μl (Gene-Tools, sequence in Figure S7) were injected followed by electroporation. Electroporation parameters were 150 V, 20 second duration, no delay, 1 Hz, train 0.5, and duration 400. Mice were assayed for splicing redirection by RT-PCR and other downstream assays 3 weeks after injection.

### *In vitro* calcium and force assays

FDB lateral and medial muscle bundles were dissected away leaving the central muscle bundle and tendon. The central muscle bundle tendon was attached to a fixed hook and the other to a force transducer. The muscle was placed in physiological saline solution, continuously gassed with 95% O_2_/5% CO_2_ at 25^°^C, and loaded with 5 μΜ Fura 4-F AM (Invitrogen). After 30 min, samples were rinsed with fresh solution and then allowed to de-esterify for 30 min. The optimal muscle length (L_o_) and voltage (V_max_) were adjusted to induce maximum twitch force. Twitch and tetanic force were measured at 1 and 150 Hz with a pulse and train durations of 0.5 and 250 ms, respectively. Fura 4-F AM excitation (360/380 nm) and emission (510 nm) were monitored simultaneously with force-frequency characteristics. After stimulation, muscle length was measured and fiber bundles were trimmed of excess muscle and connective tissue, blotted dry, and weighed. Muscle weight and L_o_ were used to estimate cross sectional area and to calculate absolute forces expressed as N/cm^2^ (59). To determine intracellular calcium changes during FDB stimulation, the 360/380 nm ratio was calculated.

### Cell Culture

C2C12 cells were maintained in DMEM with 10% FBS in 6-well tissue culture plates. To differentiate cells, C2C12 culture was grown to 100% confluency, and media was changed to DMEM with 2% horse serum. To redirect splicing in C2C12, 9-15 μΜ morpholinos were delivered by Endo-Porter (Gene-Tools) at 50% confluency. Morpholinos and Endo-Porter were added to differentiation media after cells reached 100% confluency. Differentiated C2C12 myotubes were collected at day 4.

### Immunofluorescence

Undifferentiated or differentiated C2C12 cells were grown in 6-well tissue culture plate containing glass coverslips. Cells were fixed with 4% paraformaldehyde in PBS for 15 min at room temperature. Fixed cells were washed with PBS 2x and permeabilized using 0.2% triton-X in PBS for 10 min. Cells were then blocked in 5% BSA in PBS at room temperature for 1 hour and incubated overnight in primary antibody in 5% BSA in PBS at 4^°^C. The cells were washed 3x with PBS followed by Alexa Fluor-conjugated secondary antibody incubation for 1 hour, washed 3x with PBS, DAPI stained for 5 min, and washed 3x with PBS. Deconvolution microscopy was performed by GE Healthcare Inverted Deconvolution/Image Restoration Microscope.

### Statistics

For statistical analysis, at least three samples were pooled together to determine average and variance. Error bars represent the standard deviation, and student T-test was used determine significant with P>0.05.

## Supplemental Figure Legends

**Figure S1.**
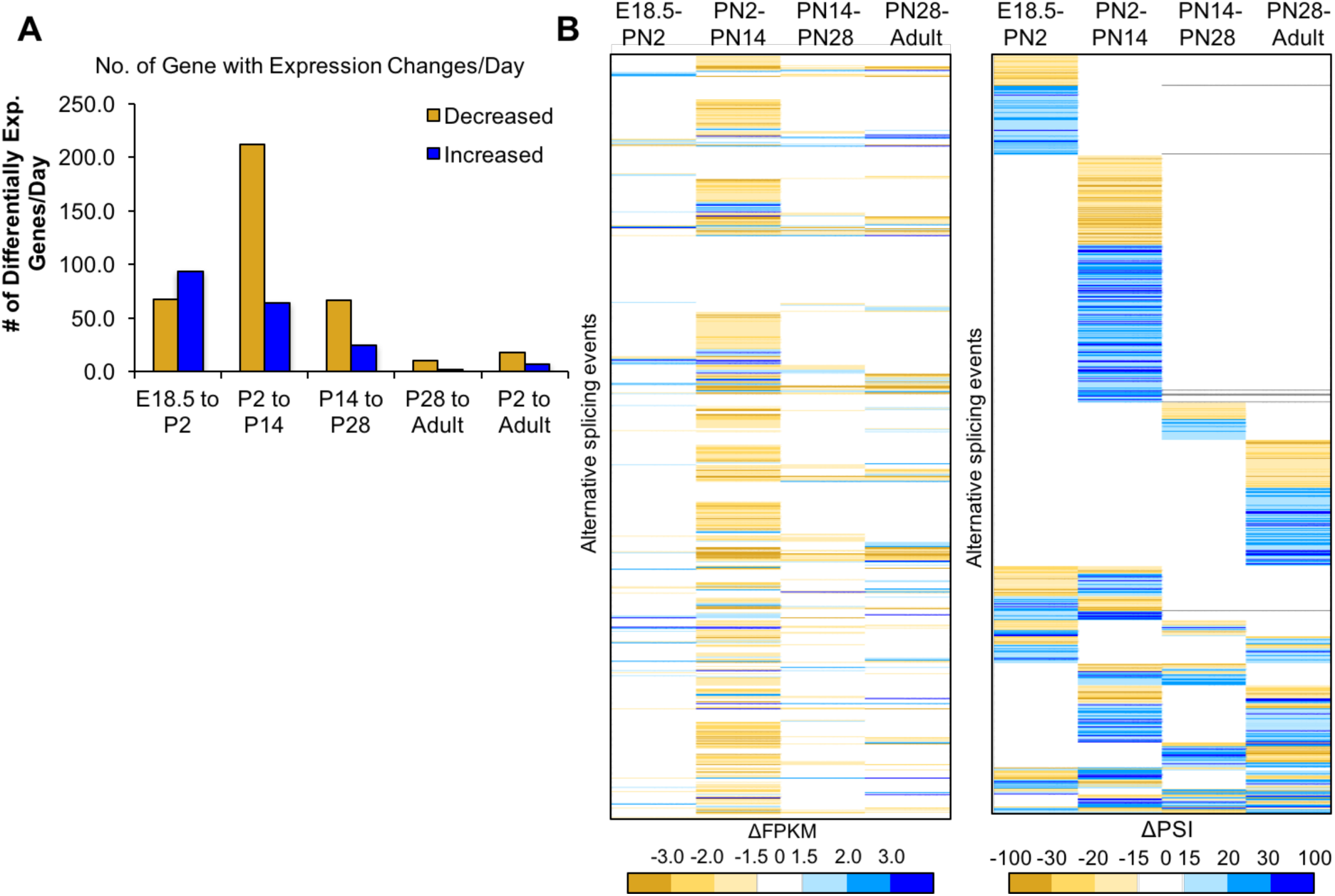
(related to Figure 2) Most changes in gene expression occur before PN14. (A) Differential gene expression changes standardized to the number of days in the time interval. (B) Heat map of differential gene expression of genes (left) with alternative splicing (right) between four intervals. The gene list is aligned in the heat maps for differential expression and alternative splicing, the latter is reproduced from Figure 2C. Largest changes are denoted by the darkest colors of red (decreased) or green (increased). White indicates unchanged.

**Figure S2.**
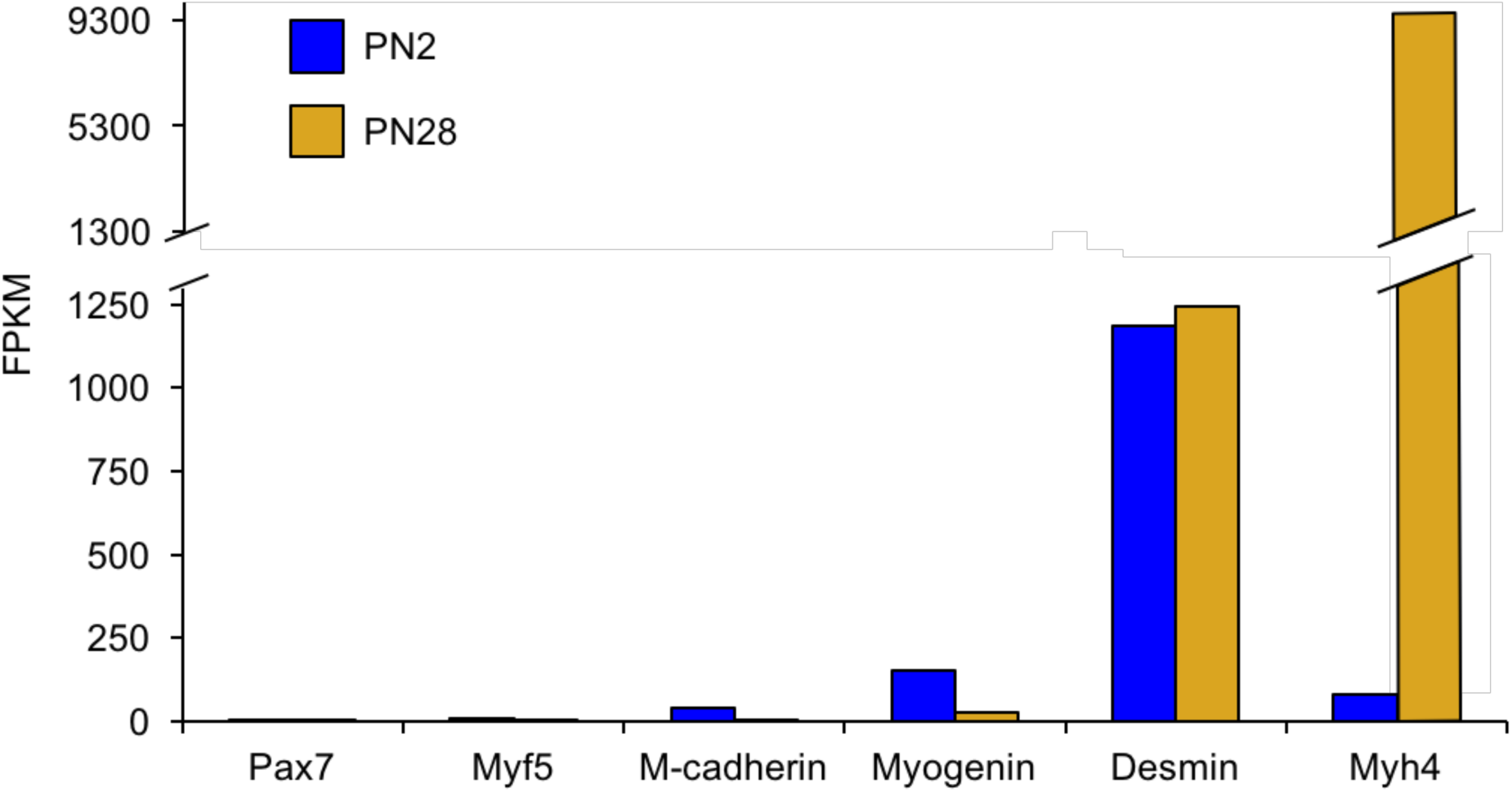
(related to Figure 1) The vast majority of transcriptome changes observed in the RNA-seq data are from myofibers with a minimal contribution from satellite cells. Genes for activated satellite cells (Pax7, Myf5, m-cadherin) had little or undetectable expression. In contrast, genes expressed in myofibers (myogenin, desmin, Myh4) are readily detected.

**Figure S3.**
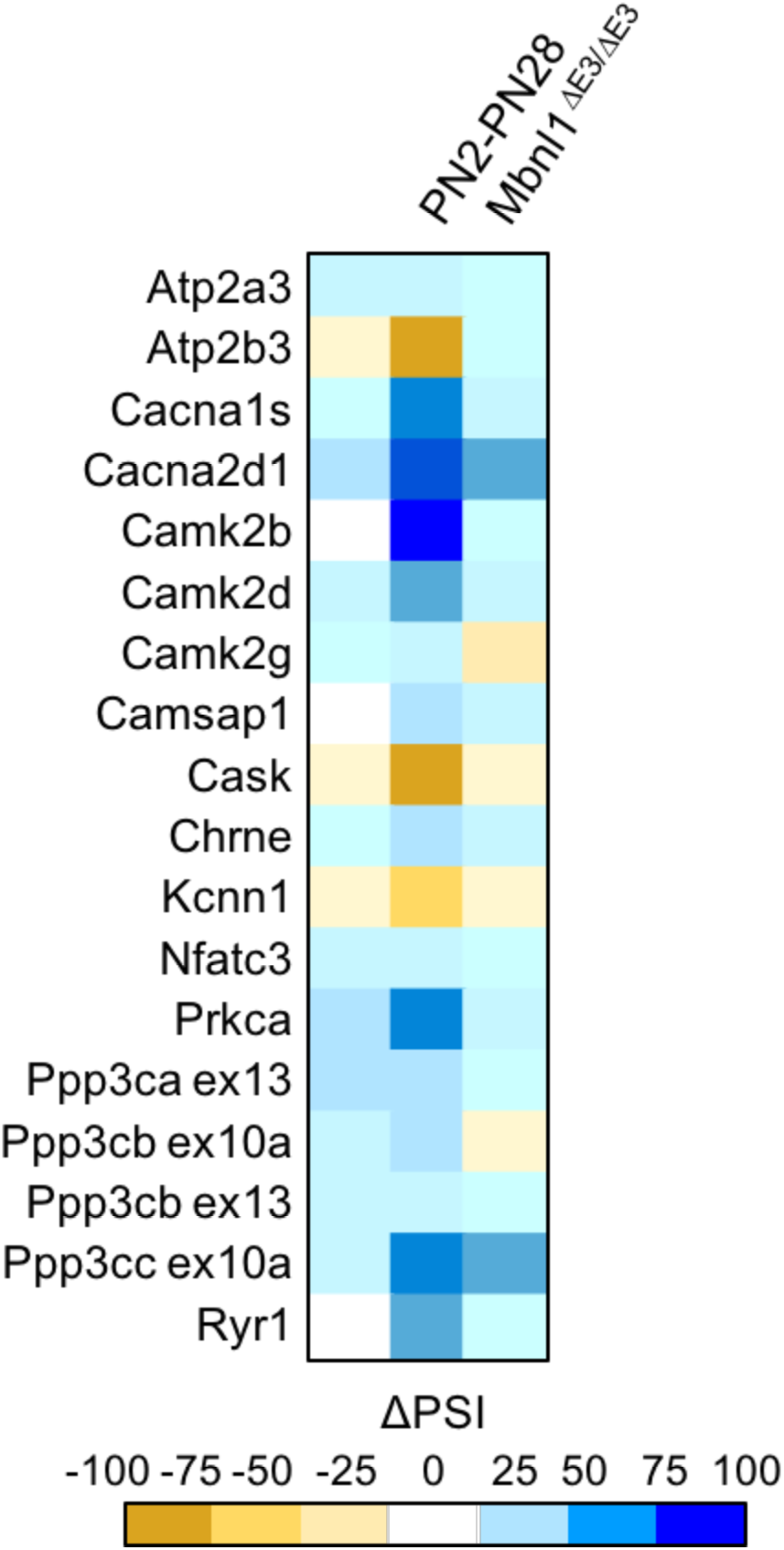
(related to Figure 6) CELF1 overexpression and Mbnl1 KO effect on calcium-handling splicing. Heat map of ΔPSI between PN2 and PN28 in wild type FVB gastrocnemius; between control mice (MDAFrtTA +dox) and hCELF1 overexpressing adult mice (MDAFrtTA/TRECUGBP1 +dox) (C57BL6/DBA;FVB) quadriceps; and between adult Mbnl1^+/+^ and Mbnl1^ΔE3/ΔE3^ (FVB) quadriceps.

**Figure S4.**
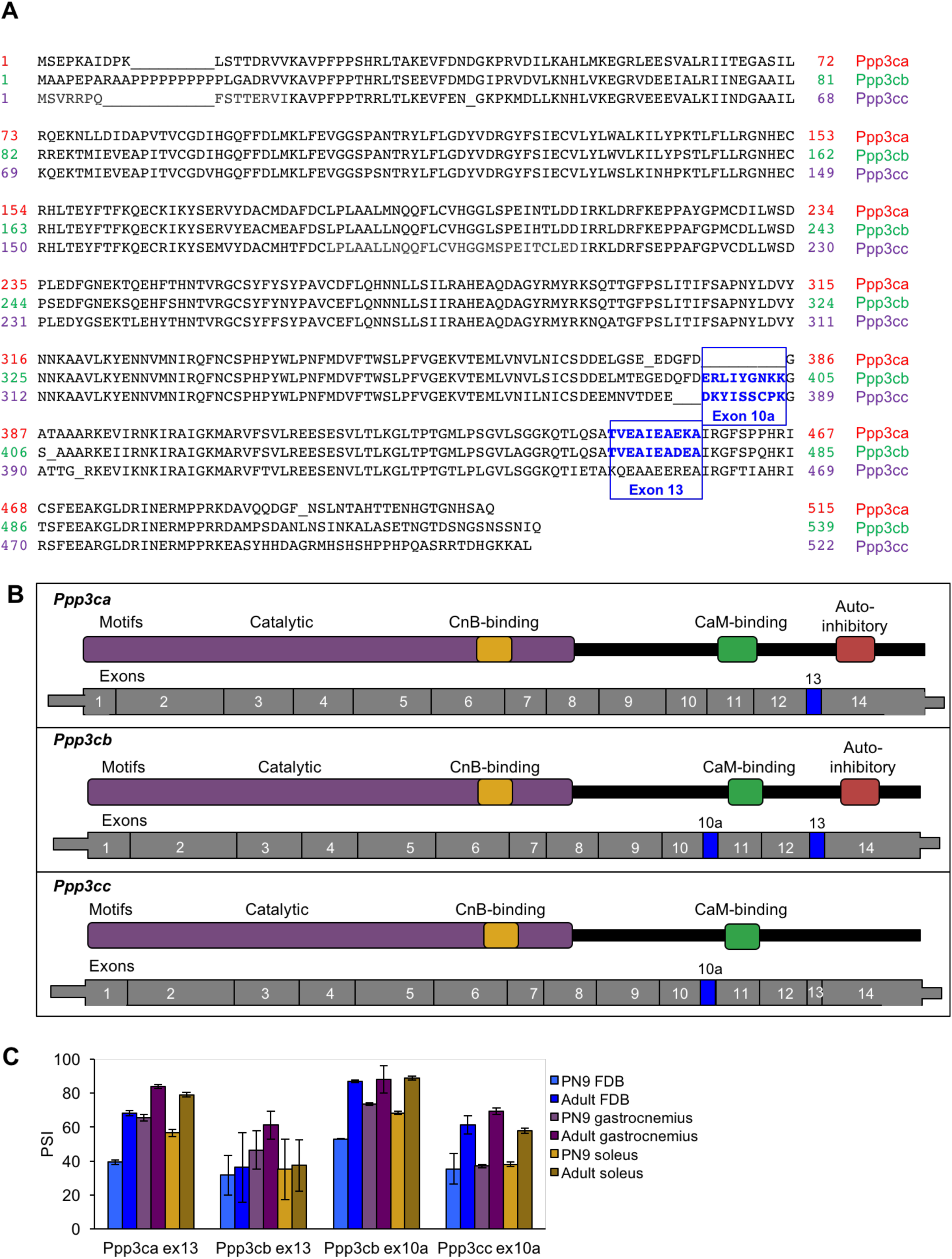
(related to Figure 7) Alignment of calcineurin A alternative exons. (A) The protein sequences of the three mouse calcineurin A paralogs are aligned. (B) Diagrams aligning calcineurin A mRNAs and proteins showing relative positions of known protein domains and alternatively spliced regions. (C) RT-PCR of calcineurin A exons during postnatal development of muscle groups with differing fiber type contributions: gastrocnemius (blue), soleus (purple), and flexor digitorum brevis (yellow). Gastrocnemius contains 6%, soleus contains 37%, and FDB contains 13% slow fiber type (type I)(60,61).

**Figure S5.**
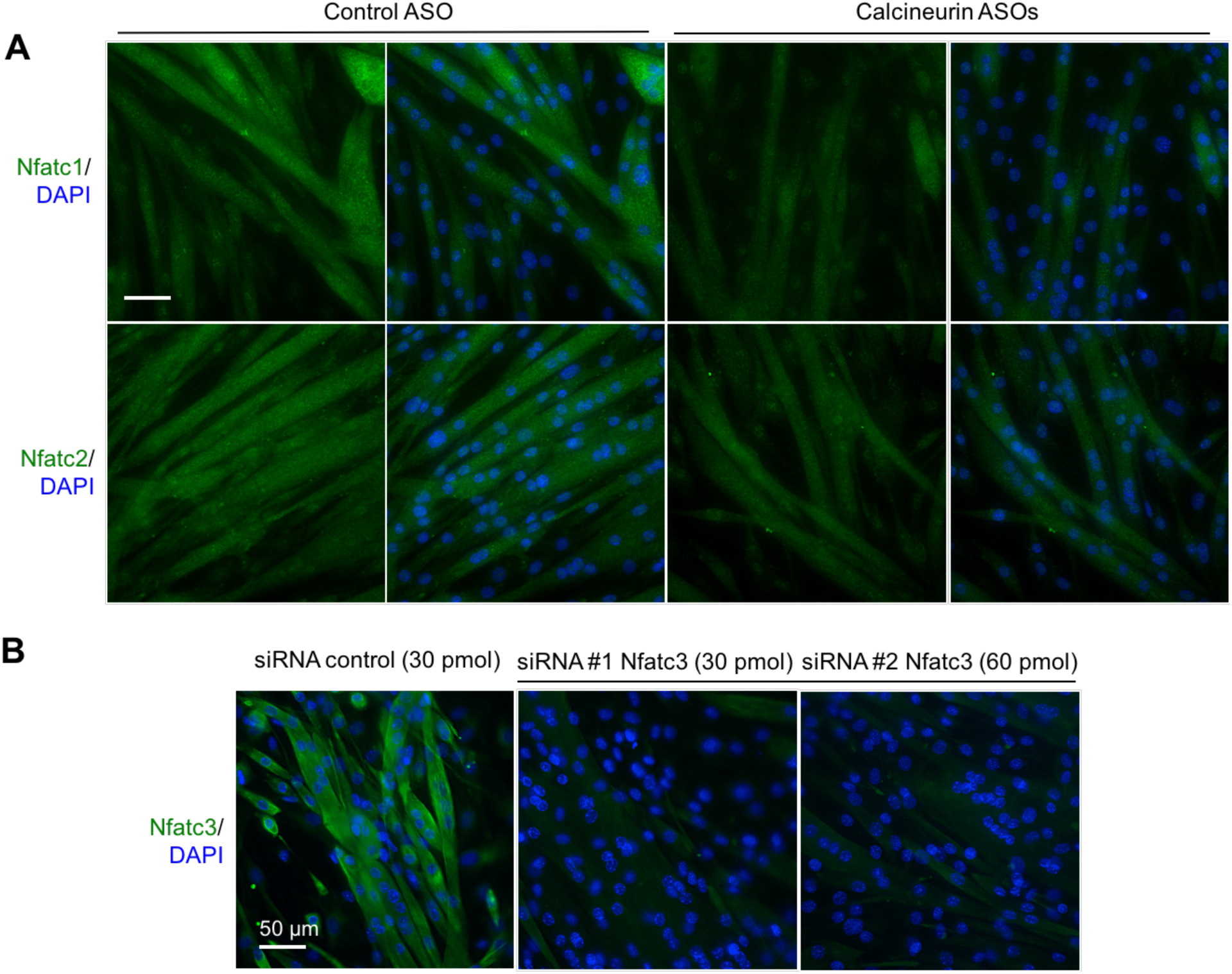
(related to Figure 8) Analysis of Nfatc1 and Nfatc2 protein localization in C2C12 and validation of Nfatc3 antibody. (A) Localization of Nfatc1 and Nfatc2 by immunofluorescence in differentiated C2C12 with ASOs. (B) Knockdown of Nfatc3 using silencer siRNAs to confirm signal of Nfatc3 antibody.

**Figure S6.**
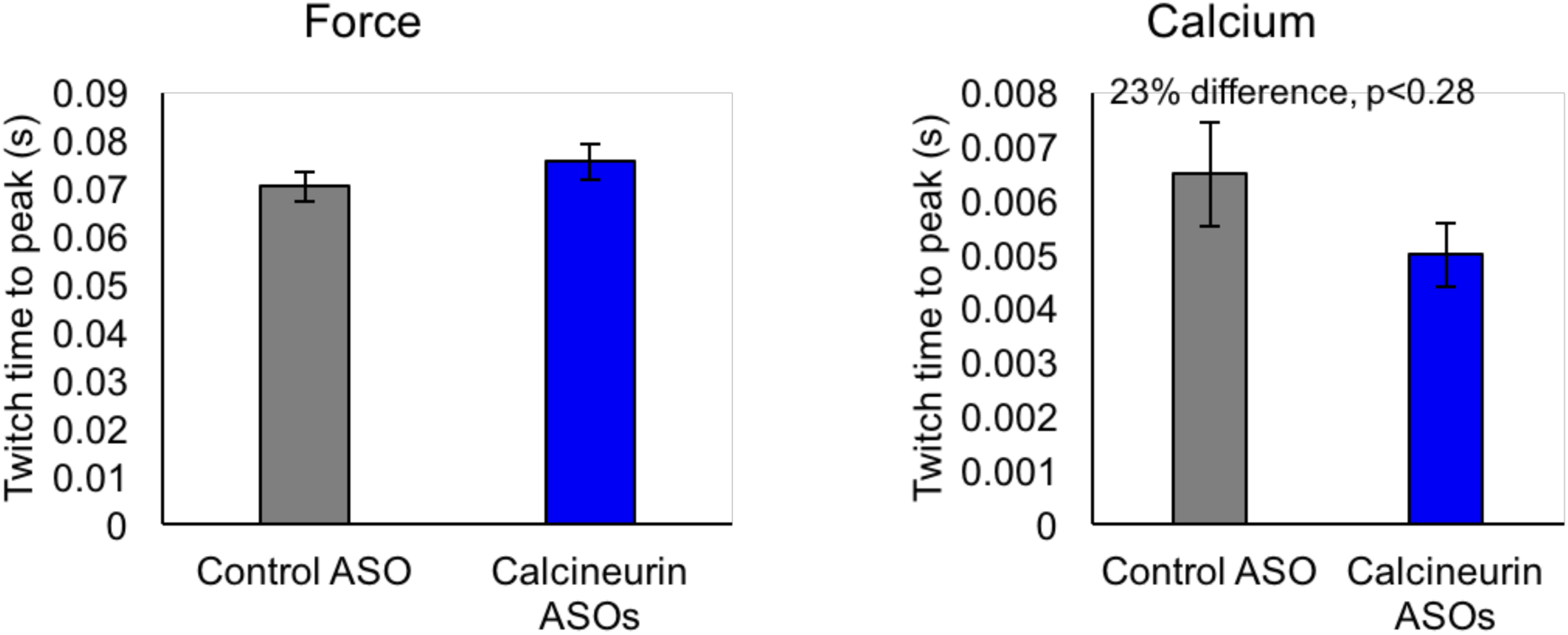
(related to Figure 9) Not significant (by student T-test) time to peak of force and calcium for control ASO and calcineurin A ASOs, n = 3-4 mice per group.

**Figure S7.**
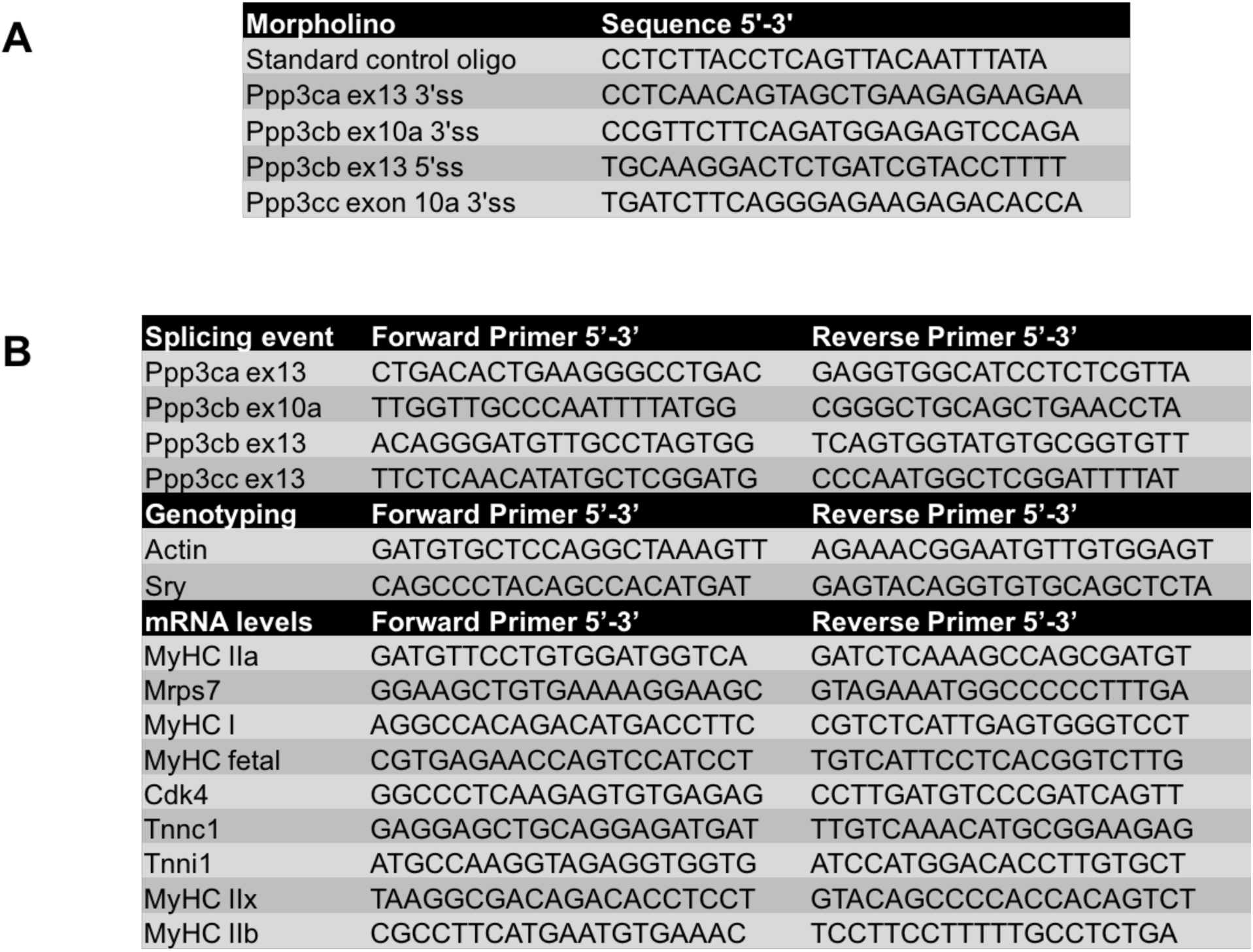
Sequences for antisense morpholinos and primers. (A) Morpholino sequences used for redirected splicing. (B) Primers used for splicing, genotyping, and mRNA levels.

## References

1. Allen RE, Merkel RA, Young RB. Cellular Aspect of Muscle Growth: Myogenic Cell Proliferation. Journal of Animal Science. The American Society of Animal Science; 1979 Jul 1;49(1):115–27.

2. Aulehla A, Pourquié O. Oscillating signaling pathways during embryonic development. Curr Opin Cell Biol. 2008 Dec;20(6): 632–7.

3. Braun T, Gautel M. Transcriptional mechanisms regulating skeletal muscle differentiation, growth and homeostasis. Nat Rev Mol Cell Biol. 2011 Jun;12(6): 349–61.

4. Buckingham M, Bajard L, Chang T, Daubas P, Hadchouel J, Meilhac S, et al. The formation of skeletal muscle: from somite to limb. J Anat. Wiley-Blackwell; 2003 Jan;202(1): 59–68.

5. Dubrulle J, Pourquié O. Coupling segmentation to axis formation. Development. 2004 Dec; 131(23):5783–93.

6. Ontell M, Feng KC, Klueber K, Dunn RF, Taylor F. Myosatellite cells, growth, and regeneration in murine dystrophic muscle: a quantitative study. Anat Rec. 1984 Feb;208(2): 159–74.

7. White RB, Biérinx A-S, Gnocchi VF, Zammit PS. Dynamics of muscle fibre growth during postnatal mouse development. BMC Dev Biol. 2010 Feb 22;10(1):21.

8. Relaix F, Zammit PS. Satellite cells are essential for skeletal muscle regeneration: the cell on the edge returns centre stage. Development. 2012 Jul 24; 139(16): 2845–56.

9. Gokhin DS, Ward SR, Bremner SN, Lieber RL. Quantitative analysis of neonatal skeletal muscle functional improvement in the mouse. J Exp Biol. 2008 Mar;211(Pt 6):837–43.

10. Franzini-Armstrong C. Simultaneous maturation of transverse tubules and sarcoplasmic reticulum during muscle differentiation in the mouse. Dev Biol. 1991 Aug;146(2): 353–63.

11. Agbulut O, Noirez P, Beaumont F, Butler-Browne G. Myosin heavy chain isoforms in postnatal muscle development of mice. Biol Cell. 2003 Sep;95(6): 399–406.

12. Gailly P, Boland B, Himpens B, Casteels R, Gillis JM. Critical evaluation of cytosolic calcium determination in resting muscle fibres from normal and dystrophic (mdx) mice. Cell Calcium. 1993 Jun;14(6): 473–83.

13. Giudice J, Xia Z, Wang ET, Scavuzzo MA, Ward AJ, Kalsotra A, et al. Alternative splicing regulates vesicular trafficking genes in cardiomyocytes during postnatal heart development. Nature Communications. Nature Publishing Group; 2014 Apr 22; 5: 3603.

14. Kalsotra A, Xiao X, Ward AJ, Castle JC, Johnson JM, Burge CB, et al. A postnatal switch of CELF and MBNL proteins reprograms alternative splicing in the developing heart. Proc Natl Acad Sci USA. 2008 Dec 23; 105(51): 20333–8.

15. Merkin J, Russell C, Chen P, Burge CB. Evolutionary dynamics of gene and isoform regulation in Mammalian tissues. Science. 2012 Dec 21;338(6114):1593–9.

16. Tress ML, Abascal F, Valencia A. Alternative Splicing May Not Be the Key to Proteome Complexity. Trends Biochem Sci. 2016 Sep 16.

17. Pickrell JK, Pai AA, Gilad Y, Pritchard JK. Noisy splicing drives mRNA isoform diversity in human cells. Dermitzakis ET, editor. PLoS Genet. 2010 Dec 9; 6(12): e1001236.

18. Gonzàlez-Porta M, Frankish A, Rung J, Harrow J, Brazma A. Transcriptome analysis of human tissues and cell lines reveals one dominant transcript per gene. Genome Biol. 2013 Jul 1;14(7):R70.

19. Barbosa-Morais NL, Irimia M, Pan Q, Xiong HY, Gueroussov S, Lee LJ, et al. The evolutionary landscape of alternative splicing in vertebrate species. Science. 2012 Dec 21;338(6114):1587–93.

20. Ellis JD, Barrios-Rodiles M, Colak R, Irimia M, Kim T, Calarco JA, et al. Tissue-specific alternative splicing remodels protein-protein interaction networks. Mol Cell. 2012 Jun 29; 46(6): 884–92.

21. Dillman AA, Hauser DN, Gibbs JR, Nalls MA, McCoy MK, Rudenko IN, et al. mRNA expression, splicing and editing in the embryonic and adult mouse cerebral cortex. Nat Neurosci. 2013 Apr;16(4): 499–506.

22. Azim S, Banday AR, Sarwar T, Tabish M. Alternatively spliced variants of gamma-subunit of muscle-type acetylcholine receptor in fetal and adult skeletal muscle of mouse. Cell Mol Neurobiol. 2012 Aug;32(6): 957–63.

23. Buck D, Hudson BD, Ottenheijm CAC, Labeit S, Granzier H. Differential splicing of the large sarcomeric protein nebulin during skeletal muscle development. J Struct Biol. 2010 May;170(2): 325–33.

24. Grande M, Suàrez E, Vicente R, Cantó C, Coma M, Tamkun MM, et al. Voltage-dependent K+ channel beta subunits in muscle: differential regulation during postnatal development and myogenesis. J Cell Physiol. Wiley Subscription Services, Inc., A Wiley Company; 2003 May;195(2): 187–93.

25. Lu S, Borst DE, Horowits R. Expression and alternative splicing of N-RAP during mouse skeletal muscle development. Cell Motil Cytoskeleton. Wiley Subscription Services, Inc., A Wiley Company; 2008 Dec;65(12): 945–54.

26. Ohsawa N, Koebis M, Suo S, Nishino I, Ishiura S. Alternative splicing of PDLIM3/ALP, for α-actinin-associated LIM protein 3, is aberrant in persons with myotonic dystrophy. Biochem Biophys Res Commun. 2011 May 27; 409(1): 64–9.

27. Sultana N, Dienes B, Benedetti A, Tuluc P, Szentesi P, Sztretye M, et al. Restricting calcium currents is required for correct fiber type specification in skeletal muscle. Development. 2016 May 1; 143(9): 1547–59.

28. Charton K, Suel L, Henriques SF, Moussu J-P, Bovolenta M, Taillepierre M, et al. Exploiting the CRISPR/Cas9 system to study alternative splicing in vivo: application to titin. Hum Mol Genet. Oxford University Press; 2016 Aug 23;:ddw280.

29. Giudice J, Loehr JA, Rodney GG, Cooper TA. Alternative Splicing of Four Trafficking Genes Regulates Myofiber Structure and Skeletal Muscle Physiology. Cell Rep. 2016 Nov 15; 17(8): 1923–33.

30. Mankodi A, Takahashi MP, Jiang H, Beck CL, Bowers WJ, Moxley RT, et al. Expanded CUG Repeats Trigger Aberrant Splicing of ClC-1 Chloride Channel Pre-mRNA and Hyperexcitability of Skeletal Muscle in Myotonic Dystrophy. Mol Cell. 2002 Jul;10(1): 35–44.

31. Charlet-B N, Savkur RS, Singh G, Philips AV, Grice EA, Cooper TA. Loss of the Muscle-Specific Chloride Channel in Type 1 Myotonic Dystrophy Due to Misregulated Alternative Splicing. Mol Cell. 2002 Jul;10(1): 45–53.

32. Lin X, Miller JW, Mankodi A, Kanadia RN, Yuan Y, Moxley RT, et al. Failure of MBNL1-dependent post-natal splicing transitions in myotonic dystrophy. Hum Mol Genet. 2006 Jul 1; 15(13): 2087–97.

33. Lee JE, Cooper TA. Pathogenic mechanisms of myotonic dystrophy. Biochem Soc Trans. Portland Press Limited; 2009 Dec;37(Pt 6):1281–6.

34. Kino Y, Washizu C, Oma Y, Onishi H, Nezu Y, Sasagawa N, et al. MBNL and CELF proteins regulate alternative splicing of the skeletal muscle chloride channel CLCN1. Nucleic Acids Res. 2009 Oct;37(19): 6477–90.

35. Wang ET, Ward AJ, Cherone JM, Giudice J, Wang TT, Treacy DJ, et al. Antagonistic regulation of mRNA expression and splicing by CELF and MBNL proteins. Genome Res. Cold Spring Harbor Lab; 2015 Jun;25(6): 858–71.

36. Konieczny P, Stepniak-Konieczna E, Sobczak K. MBNL proteins and their target RNAs, interaction and splicing regulation. Nucleic Acids Res. Oxford University Press; 2014; 42(17): 10873–87.

37. Dasgupta T, Ladd AN. The importance of CELF control: molecular and biological roles of the CUG-BP, Elav-like family of RNA-binding proteins. Wiley Interdisciplinary Reviews: RNA. John Wiley & Sons, Inc; 2012 Jan 1;3(1):104–21.

38. Ward AJ, Rimer M, Killian JM, Dowling JJ, Cooper TA. CUGBP1 overexpression in mouse skeletal muscle reproduces features of myotonic dystrophy type 1. Hum Mol Genet. Oxford University Press; 2010 Sep 15; 19(18): 3614–22.

39. Kanadia RN, Johnstone KA, Mankodi A, Lungu C, Thornton CA, Esson D, et al. A muscleblind knockout model for myotonic dystrophy. Science. 2003 Dec 12;302(5652):1978–80.

40. Al-Shanti N, Stewart CE. Ca 2+/calmodulin-dependent transcriptional pathways: potential mediators of skeletal muscle growth and development. Biological Reviews. Blackwell Publishing Ltd; 2009 Nov;84(4): 637–52.

41. Ip JY, Tong A, Pan Q, Topp JD, Blencowe BJ, Lynch KW. Global analysis of alternative splicing during T-cell activation. RNA. Cold Spring Harbor Lab; 2007 Apr;13(4): 563–72.

42. Singh RK, Xia Z, Bland CS, Kalsotra A, Scavuzzo MA, Curk T, et al. Rbfox2-coordinated alternative splicing of Mef2d and Rock2 controls myoblast fusion during myogenesis. Mol Cell. 2014 Aug 21; 55(4): 592–603.

43. Trapnell C, Williams BA, Pertea G, Mortazavi A, Kwan G, van Baren MJ, et al. Transcript assembly and quantification by RNA-Seq reveals unannotated transcripts and isoform switching during cell differentiation. Nat Biotechnol. Nature Research; 2010 May;28(5): 511–5.

44. Kimura T, Pace SM, Wei L, Beard NA, Dirksen RT, Dulhunty AF. A variably spliced region in the type 1 ryanodine receptor may participate in an inter-domain interaction. Biochem J. Portland Press Limited; 2007 Jan 1;401(1):317–24.

45. Kimura T, Lueck JD, Harvey PJ, Pace SM, Ikemoto N, Casarotto MG, et al. Alternative splicing of RyR1 alters the efficacy of skeletal EC coupling. Cell Calcium. 2009 Mar;45(3): 264–74.

46. Periasamy M, Kalyanasundaram A. SERCA pump isoforms: their role in calcium transport and disease. Muscle Nerve. Wiley Subscription Services, Inc., A Wiley Company; 2007 Apr;35(4): 430–42.

47. Delling U, Tureckova J, Lim HW, De Windt LJ, Rotwein P, Molkentin JD. A calcineurin-NFATc3-dependent pathway regulates skeletal muscle differentiation and slow myosin heavy-chain expression. Mol Cell Biol. American Society for Microbiology (ASM); 2000 Sep;20(17): 6600–11.

48. Buljan M, Chalancon G, Eustermann S, Wagner GP, Fuxreiter M, Bateman A, et al. Tissue-specific splicing of disordered segments that embed binding motifs rewires protein interaction networks. Mol Cell. 2012 Jun 29; 46(6): 871–83.

49. Felkin LE, Narita T, Germack R, Shintani Y, Takahashi K, Sarathchandra P, et al. Calcineurin splicing variant calcineurin Aβ1 improves cardiac function after myocardial infarction without inducing hypertrophy. Circulation. American Heart Association, Inc; 2011 Jun 21; 123(24): 2838–47.

50. Gómez-Salinero JM, López-Olañeta MM, Ortiz-Sánchez P, Larrasa-Alonso J, Gatto A, Felkin LE, et al. The Calcineurin Variant CnAβ1 Controls Mouse Embryonic Stem Cell Differentiation by Directing mTORC2 Membrane Localization and Activation. Cell Chem Biol. 2016 Nov 17; 23(11): 1372–82.

51. Giudice J, Xia Z, Li W, Cooper TA. Neonatal cardiac dysfunction and transcriptome changes caused by the absence of Celf1. Sci Rep. Nature Publishing Group; 2016 Oct 19; 6: 35550.

52. Orengo JP, Ward AJ, Cooper TA. Alternative splicing dysregulation secondary to skeletal muscle regeneration. Ann Neurol. Wiley Subscription Services, Inc., A Wiley Company; 2011 Apr;69(4): 681–90.

53. Han H, Irimia M, Ross PJ, Sung H-K, Alipanahi B, David L, et al. MBNL proteins repress ES-cell-specific alternative splicing and reprogramming. Nature. Nature Research; 2013 Jun 13;498(7453):241–5.

54. Kroeze Y, Oti M, van Beusekom E, Cooijmans RHM, van Bokhoven H, Kolk SM, et al. Transcriptome Analysis Identifies Multifaceted Regulatory Mechanisms Dictating a Genetic Switch from Neuronal Network Establishment to Maintenance During Postnatal Prefrontal Cortex Development. Cereb Cortex. Oxford University Press; 2017 Jan 19.

55. Vernia S, Edwards YJ, Han MS, Cavanagh-Kyros J, Barrett T, Kim JK, et al. An alternative splicing program promotes adipose tissue thermogenesis. Elife. eLife Sciences Publications Limited; 2016 Sep 16; 5: 1986.

56. Trapnell C, Pachter L, Salzberg SL. TopHat: discovering splice junctions with RNA-Seq. Bioinformatics. 2009 Apr 23; 25(9): 1105–11.

57. Robinson MD, McCarthy DJ, Smyth GK. edgeR: a Bioconductor package for differential expression analysis of digital gene expression data. Bioinformatics. 2010 Nov 11; 26(1): 139–40.

58. Katz Y, Wang ET, Airoldi EM, Burge CB. Analysis and design of RNA sequencing experiments for identifying isoform regulation. Nature Methods. Nature Research; 2010 Nov 7; 7(12): 1009–15.

59. Close RI. Dynamic properties of mammalian skeletal muscles. Physiological Reviews. American Physiological Society; 1972 Jan;52(1): 129–97.

60. Augusto V, Padovani CR, Campos G. Skeletal muscle fiber types in C57BL6J mice. Braz J Morphol Sci. 2004.

61. González E, Messi ML, Zheng Z, Delbono O. Insulin-like growth factor-1 prevents age-related decrease in specific force and intracellular Ca2+ in single intact muscle fibres from transgenic mice. The Journal of Physiology. Blackwell Publishing Ltd; 2003 Nov 1; 552(3): 833–44.

